# Extracellular Caspase-1 regulates hair follicle stem cell migration during wound-healing

**DOI:** 10.1101/548529

**Authors:** Subhasri Ghosh, Akshay Hegde, Akhil SHP Ananthan, Sunny Kataria, Neha Pincha, Anupam Dutta, Abhik Dutta, Srilekha Athreya, Sneha Khedkar, Rakesh Dey, Aishwarya Bhosale, Colin Jamora

**Affiliations:** IFOM-inStem Joint Research Laboratory, Centre for Inflammation and Tissue Homeostasis, Institute for Stem Cell Science and Regenerative Medicine, Bangalore, Karnataka, India; School of Chemical and Biotechnology (SCBT), Shanmugha Arts, Science, Technology and Research Academy (SASTRA), deemed to be University, Thanjavur 613401, Tamil Nadu, India; School of Dentistry, University of California, San Francisco; Cologne Excellence Cluster for Stress Responses in Ageing-associated Diseases (CECAD), Cologne, 50931, Germany

**Author notes:** indicates equal contribution (Subhasri Ghosh, Akshay Hegde, Akhil SHP Ananthan).

## Abstract

The wound-healing process is a paradigm of the directed migration of various pools of stem cells from their niche to the site of injury where they replenish damaged cells. Two decades have elapsed since the observation that wounding activates multipotent hair follicle stem cells to infiltrate the epidermis, but the cues that coax these cells out of their niche remains unknown. Using an excisional wound and genetic mouse models of wound healing, we discovered that Caspase-1, a protein classically known as an integral component of the cytosolic inflammasome, is secreted and has a non-canonical role in the extracellular milieu. Through its Caspase Activation Recruitment Domain (CARD), Caspase-1 is sufficient to initiate chemotaxis of hair follicle stem cells into the epidermis. Uncovering this novel function of Caspase-1 also facilitates a deeper understanding of the mechanistic basis of the epithelial hyperplasia found to accompany numerous inflammatory skin diseases.

## INTRODUCTION

Migration of stem cells from one niche to another is a fundamental behavior observed during tissue morphogenesis, homeostasis, and repair. A common thread running throughout these phenomena is the ability of stem cells to sense their environmental cues that, in turn, regulate their spatiotemporal localization with amazing precision. Upon arriving at their destination, these stem cells differentiate to form, regenerate, or repair a tissue (Goodell et al., 2015; Laird et al., 2008). Perturbations of such cellular responses underlie a spectrum of pathologies ranging from developmental defects, tumor metastasis and ineffective wound closure (Ge and Fuchs, 2018). In somatic tissues, the wound-healing process in the skin is a paradigm of the directed migration of various cutaneous stem cell pools to the site of injury where they differentiate to replenish lost or damaged cells (Gonzales and Fuchs, 2017; Rognoni and Watt, 2018). While there has been substantial investment and progress in understanding the lineage trajectory of stem cells once they reach their destination, comparatively little is understood regarding the mechanisms guiding their chemotactic journey to the wound site. As the outermost protective barrier of our body, the skin has evolved the ability to heal itself rapidly owing to proliferation and migration of multiple stem cell pools in various cutaneous niches into the wound microenvironment (Gonzales and Fuchs, 2017; Plikus et al., 2012). The process of wound-healing occurs in three temporally overlapping phases; the inflammatory, the proliferative and remodeling phases, each of which is characterized by extensive cellular crosstalk, leading to repair of the damaged tissue (Gurtner et al., 2008; Rodrigues et al., 2019). Briefly, the inflammatory phase is marked by immune cell infiltration into the wound-bed and release of cytokines and growth factors. This creates a foundation to support the ensuing proliferative phase during which neighboring stem cells proliferate and migrate into the wound to aid in rapid tissue formation and wound closure (Gonzales and Fuchs, 2017; Plikus et al., 2012). Reformation of the epidermal barrier by proliferation and migration of keratinocytes, known as re-epithelialization, is enhanced by the contribution of hair follicle stem cells (HFSCs) differentiating to epidermal keratinocytes, an emergency response not observed otherwise in homeostasis (Dekoninck and Blanpain, 2019). Though wound directed migration of HFSCs had been first observed more than a decade ago, the guidance cues have remained elusive (Ito et al., 2005; Tumbar et al., 2004). It is becoming increasingly clear that inflammatory signals from resident and recruited immune cells are a major regulator of HFSC behavior in both homeostasis and wound-healing (Naik et al., 2017). Interestingly this immune cell - stem cell crosstalk is emerging as a common theme for the regulation of epidermal stem cells in general (Kobayashi et al., 2019; Mathur et al., 2019; Naik et al., 2018). Therefore, identification of an unexpected migratory stimulus arising from an inflammatory signaling cascade is relevant not only in the context of wound healing but also skin carcinomas where physiological response mechanisms of epithelial stem cells are usurped (Rognoni and Watt, 2018; Sundaram et al., 2018). Our findings reveal a novel function of a common inflammatory molecule in directed cell migration.

## RESULTS

### Migration of hair follicle stem cells (HFSCs) into the epidermis is recapitulated in the caspase-8 conditional knockout model of wound healing

A major hurdle in dissecting the mechanisms regulating the homing of HFSCs to the wound bed has been the limited number of responsive HFSCs. Only the hair follicles that are immediately adjacent to the wound activate their stem cells in response to the damage stimuli (Ito et al., 2005). A genetic model of wound-healing that elicits a repair process throughout the skin in the absence of an injury would bypass this technical obstacle by amplifying the extensive intercellular crosstalk that takes place during the tissue regeneration program. Thus, we evaluated the potential of studying the mechanisms guiding HFSC migration with the aid of a mouse with the conditional knock-out of epidermal caspase-8 (C8cKO), which we previously reported to mimic many phenomena associated with the inflammatory, proliferative and remodeling phases of the wound healing program (Lee et al., 2015, 2009, 2017). Mechanistic insights derived from this mouse model were then verified using excisional wounds on the wild-type mouse skin.

HFSCs and their progeny have been reported to robustly occupy the epidermis between 2 – 5 days following wounding (Adam et al., 2015; Ito et al., 2005). A priori, one would expect that stem cell mobilization would be initiated soon after injury given their critical role in rapid wound repair. Surprisingly, whether changes in HFSC activity can be observed at earlier timepoints post-wounding have not been reported. In order to fill this gap in our understanding of the initial cellular response of HFSCs to tissue damage, we examined the behavior of these cells in the 8 – 24 hour time-frame, a period when even proliferation in the HFSC niche has yet to begin (Figure 1-figure supplement 1A). The localization of HFSCs in the mouse skin has been studied using genetic lineage reporter models (Adam et al., 2015; Ito et al., 2005; Kretzschmar and Watt, 2012; Morris et al., 2004) or by mapping endogenous markers such as Sox9, a master regulator of HFSC identity. Sox9 expression has been used extensively to trace HFSCs in development (Nowak et al., 2008; Vidal et al., 2005), wound-healing (Adam et al., 2015; Ge et al., 2017; Kasper et al., 2011) and in pathological transformation (Larsimont et al., 2015; Shi et al., 2013; Wong and Reiter, 2011). With this bona-fide endogenous marker of HFSCs we have studied their spatio-temporal localization in homeostatic and wounded skin as well as the genetic model of wound healing. As previously reported (Kadaja et al., 2014; Vidal et al., 2005), under homeostatic conditions in adult mice at the telogen stage of the hair cycle, HFSCs expressing Sox9 are clustered at the base of the hair follicle in their niche known as the bulge (Figure 1A). However, upon a full thickness excisional wound, as early as 16 hours post-wounding, there is a noticeable amount of HFSCs migration upwards from the bulge along the outer layer of the hair follicles towards the infundibulum (the upper portion of the hair follicle, adjacent to the epidermis) (Figure 1A). By 24 hours post wounding, there is a substantial amount of HFSCs in the infundibulum area and some infiltration of these cells into the epidermis (Figure 1A). Experiments where a lineage reporting strategy has been used to trace HFSCs upon wounding revealed their upward migration towards the damaged epidermis within 24 hours (Tumbar et al., 2004).

**Figure 1:**
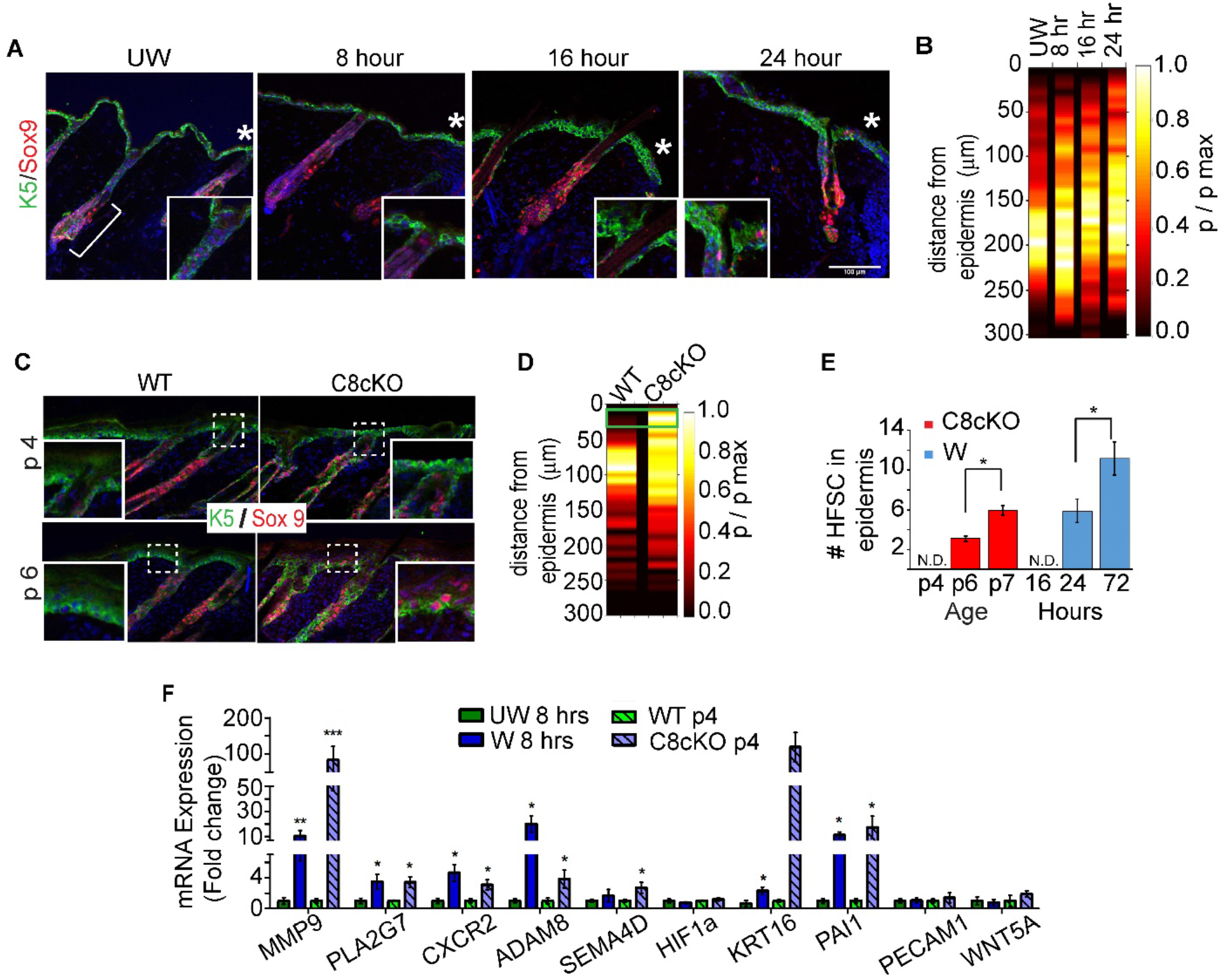
Epidermal knock-out of Caspase-8 (C8cKO) recapitulates HFSC migration to the epidermis. **(A)** Immunofluorescence images of localization of HFSCs marked by Sox9 (red) in adult skin where the hair follicle and epidermis are marked by keratin-5 (green) in unwounded (UW) and post-wounded skin. Bracket denotes the bulge; and boxed region marks the infundibulum (magnified in insets). “*” marks the wound edge Scale bar = 100um. (**B**) Probability distribution of HFSCs along the hair follicle in unwounded (UW) and wounded (W) skin at 8, 16 and 24 hours post wounding, represented as a heat-map with distance from the epidermis represented on the y-axis (see Figure 1 Supplement 2B and Supplementary table 1). (**C**) HFSC localization in the skin of wild-type (WT) and caspase-8 conditional knock-out (C8cKO) mice at postnatal days 4 (p4) and 6 (p6). Inset showing infundibulum at p4 and interfollicular epidermis at p6. (**D**) Probability distribution of HFSCs along hair follicles in the skin of WT and C8cKO mice at p4. Green box marks the infundibulum region of hair follicles (see Figure 1: Figure Supplement 2E and Supplementary table 3). (**E**) HFSC numbers analysed from the interfollicular epidermis (50 by 50 mm^2^ regions) in C8cKO mice at postnatal days 4, 6 and 7 (red bars); and compared to the wound edge epidermis at 16, 24 and 72 hours of wound-healing (W) (blue bars). Mean and S.E.M is calculated from 5 biological replicate. **(F)** Expression analysis of HFSC specific wound activated migration genes from unwounded (UW, green) and wounded skin (W, blue) at 8 hours of wound-healing, compared to WT (green hatch) and C8cKO (blue hatch) p4 epidermis and hair follicles. Data for 8 hrs timepoints is a composite of 6 biological replicates and 3 technical replicates, and for p4 timepoints 5 biological replicates and 3 technical replicates Error bars represent S.E.M. “*” indicates p<0.05, “**” indicates p<0.01, “***” indicates p<0.001 “n.s.” indicates “not significant”.

To quantitatively track the changes in HFSC localization upon wounding, we developed an image analysis algorithm which measures the distance of individual HFSCs from the junction between the epidermis and the hair follicle (Figure 1-figure supplement 2A), and calculates their frequency of occurrence in bins of equal dimensions along the length of the hair follicle (Figure 1: figure supplement 2B). As the response of HFSCs to wounding occurs immediately adjacent to the wound edge, with the response waning further away from the wound margin (Ito et al., 2005; Jaks et al., 2008), we have focused our analysis on the first hair follicle adjacent to the wound bed to control for uniformity in the different experimental backgrounds (Figure 1-figure supplement 1B). In Figure 1B, we represent the quantitative distribution of HFSCs along hair follicles as a heatmap, where the regions having larger numbers of HFSCs are represented in lighter colors. Consistent with both published reports (Kadaja et al., 2014; Nowak et al., 2008; Vidal et al., 2005) and with the qualitative localization of Sox9 expressing HFSCs, the highest probability of occurrence of HFSCs is detected in the bulge (∼ 150 - 250 μm from the epidermis) of hair-follicles in both unwounded and wounded skin. We did not observe a statistically significant increase in the relocalization of the HFSCs near the infundibulum (within 50 μm from the epidermis) at 8 hours after wounding (Figure 1-figure supplement 2B). However, by 16 hours post-wounding, we observed a measurable increase in the frequency of HFSCs close to the infundibulum and by 24 hours, they are highly represented in the infundibulum (Figure 1-figure supplement 2B). Thus we have uncovered a novel phenomenon wherein migration of HFSCs precedes their proliferation in the niche, typically observed in the bulge after 72 hours of wound-healing (Mardaryev et al., 2011). This observation is similar to the behavior of keratinocytes in the wounded epidermis with distinct zones -the immediate wound-proximal region of the epidermis contains only migrating cells, which is followed by a separate zone of proliferating keratinocytes (Aragona et al., 2017; Park et al., 2017). Our observation that emigration of HFSCs from the bulge precedes proliferation implies that the exodus of stem cells may be a signal to the stationary HFSCs to multiply and repopulate the depleted niche.

To test if HFSC migration in response to injury is recapitulated in the genetic model of wound-healing (C8cKO), we knocked-out caspase-8 expression in the epidermis of mice using the constitutively active K14-Cre driver. We observed a noticeable accrual of HFSCs localization within the infundibulum region of the hair follicle (< 50μm from the epidermis) (Figure 1C) Similar to HFSCs in wound-proximal hair follicles, the HFSCs in the C8cKO mouse exhibits migration into the epidermis above the site of arrector pilli muscle (APM) attachment demarcating the HFSC niche (Figure 1-figure supplement 2D). Using the image analysis algorithm, we have observed a statistically significant increase in the HFSC localization within the infundibulum of the postnatal day 4 (p4) C8cKO skin (Figure 1D, Figure 1-figure supplement 2E). Interestingly, at this p4 timepoint, in the genetic model of wounding, we do not detect any Sox9 expressing HFSCs in the epidermis (Figure 1E). Nevertheless, as the wound healing phenotype advances in postnatal day 6 mice, there is a progressive increase in the number of HFSCs detected in the interfollicular epidermis (Figures 1C and E). This result recapitulates the phenomenon observed in the edge of an excisional wound wherein there is increasing amounts of HFSCs in the epidermis beginning one day after wounding (Figures 1E). To test if adult mice recapitulate the Sox9 migration phenomenon observed in the neonatal (p4) mice, we knocked-out caspase-8 in the adult mice using tamoxifen inducible Cre expression driven by the K14 promoter. In the adult mice, Sox9 cells are localized in the bulge region of the hair follicle, but two weeks post tamoxifen induction, Sox9 cells are found in the infundibulum region of the hair follicles of the adult C8cKO mice (Figure 1 - figure supplement 3). Thus, the genetic model of wound-healing can elicit epidermal homing of the HFSCs, where they contribute to the epidermal hyperplasia phenotype of the C8cKO mouse (Lee et al., 2009).

To further confirm the trend of HFSCs redistribution towards the epidermis as a chemotactic process, we scored for the activation of genes that can impact their migratory behavior. Analysis of published transcriptome data obtained from HFSCs isolated from unwounded and wounded skin (Ge et al., 2017) (Figure 1-figure supplement 4A) yielded a set of genes ontologically associated with cell migration (Figure 1-figure supplement 4B). From this dataset, selected migration-associated genes were tested for their expression at different time points post excisional wounding (Figure 1-figure supplement 4C). This analysis demonstrated a significant upregulation of migratory genes by 8 hours following excisional wounding (Figure 1F). The observation that the HFSC migration gene program is activated (at 8 hours) in excisional wounds even before a measurable HFSC flux out of the niche (at 16 hours), supports previous observations that HFSCs activate the wound-response program at the gene expression level considerably before they physically start migrating out of their niche (Joost et al., 2018). A similar trend in the expression of the migration genes is seen in the epithelial compartment of the C8cKO skin from p4 mice prior to their entrance into the interfollicular epidermis of the skin (Figure 1F). Altogether, these results suggest that the C8cKO mouse recapitulates both the migratory gene signature as well as the motility behavior of the HFSCs in response to an excisional wound. We, therefore, exploited the C8cKO skin as a platform to dissect the molecular mechanisms regulating the homing behavior of HFSCs from the hair follicle to the epidermis.

### Caspase-1 is a critical regulator of HFSC migration to the epidermis

Unlike the excisional wound model in which the HFSCs are mobilized only within in the follicles adjacent to the wound bed, the advantage of the C8cKO model is the activation of HFSCs in follicles uniformly throughout the skin. We thus used this genetic model to gain clues about the mechanism underlying the chemotactic behavior of HFSCs. An important feature that is conserved between the excisional wound and the C8cKO skin is the activation of the Caspase-1 containing inflammasome. The inflammasome is a macromolecular complex in the cytosol that mediates the unconventional secretion of many inflammatory cytokines including IL-1α (Lee et al., 2015, 2009). In particular, IL-1α mediates the reciprocal interactions between epidermal keratinocytes and dermal fibroblasts to promote proliferation of basal epidermal stem cells. Moreover, IL-1α also mediates the crosstalk between epidermal keratinocytes and resident dendritic epidermal T-cells to promote HFSC proliferation (Lee et al., 2017). Consequently, caspase-1 null mice (C1^-/-^) exhibit a delay in wound closure by affecting epithelial stem cell proliferation in different niches of the skin (Lee et al., 2015). We observed that caspase-1 expression is upregulated as early as 8 hours post wounding (Figure 2-figure supplement 1A), a timeframe corresponding with the upregulation of HFSC migration genes. Given its effects on proliferation, we hypothesized that caspase-1 may also influence re-epithelialization of the wound by regulating the migration of HFSCs to the epidermis. To test this hypothesis, we analyzed the distribution of HFSCs along the hair follicles in C8cKO skin lacking caspase-1 (C8cKO/C1^-/-^) and compared it to their localization in C8cKO and WT skins (Figure 2A). The HFSCs found to be epidermally directed (present within the infundibulum) in the C8cKO skin at p4, is substantially reduced in the C8cKO/C1^-/-^ skin (Figure 2A). Upon confirming that the localization of the HFSC niche is consistent among these three genetic backgrounds (Figure 2-figure supplement 1B), we performed quantitative analysis of HFSC localization along hair follicles in these mice. We found that the increased frequency of HFSCs in the infundibulum in the C8cKO mice relative to its wild-type counterpart is significantly reduced in the absence of caspase-1 (C8cKO/C1^-/-^) (Figure 2B, Figure 2-figure supplement 1C). Similar to the hair follicles in the neonatal mice, the absence of caspase-1 in the adult C8cKO mice also exhibited reduced epidermal directionality of HFSCs (Figure 2-figure supplement 2). This suggests that the initial chemotaxis of HFSCs towards the epidermis is largely dependent on caspase-1.

**Figure 2:**
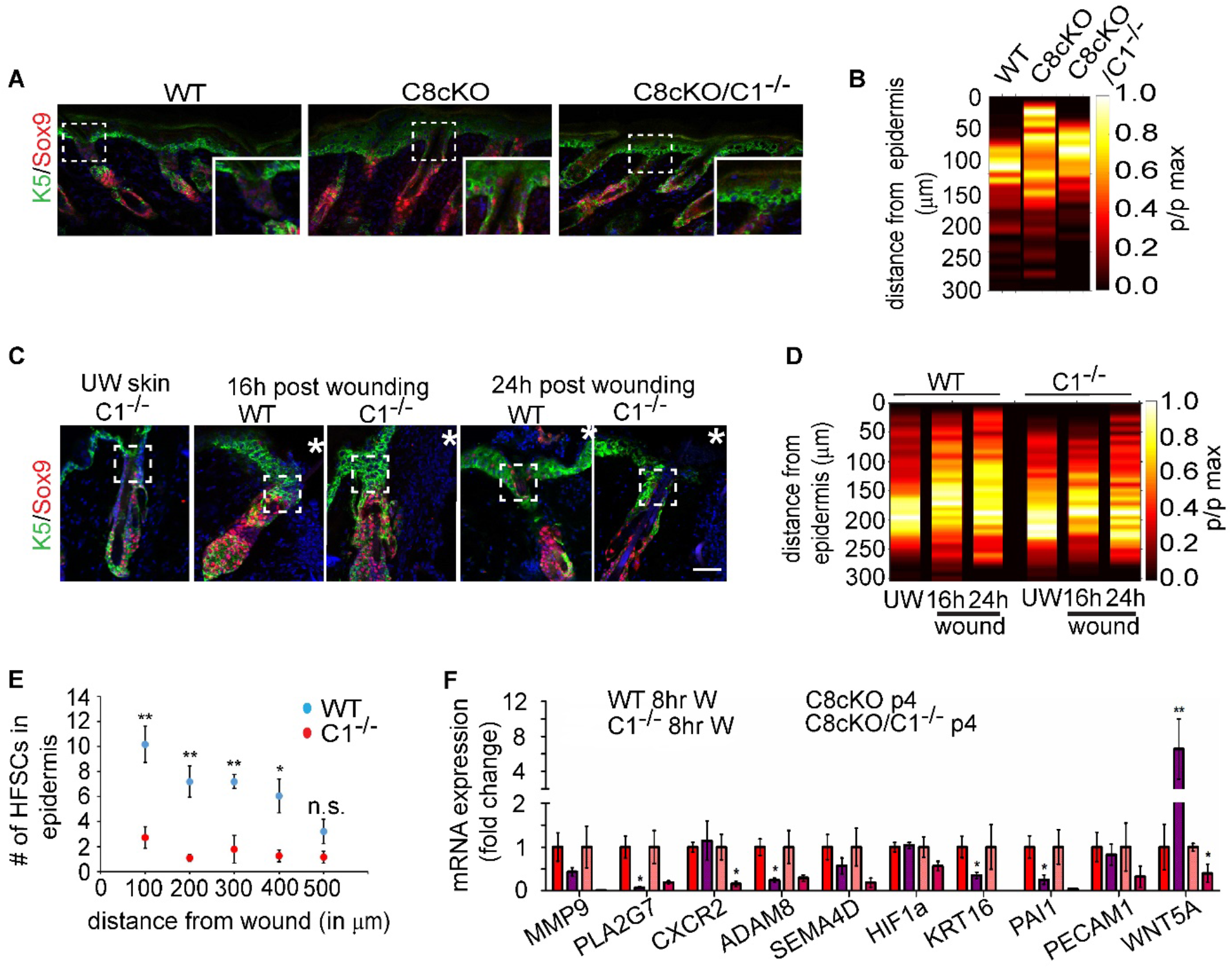
Caspase-1 regulates HFSC migration to the epidermis. **(A)** Immunofluorescence images of localization of HFSCs (Sox9, red) in WT, C8cKO and C8cKO/C1^-/-^ skin at postnatal day 4. Epidermis and hair follicles are marked by K5 (green) Insets show the infundibulum of representative hair follicles. (**B**) Probability distribution of HFSCs along hair follicles analyzed from WT, C8cKO and C8cKO/C1^-/-^ skin at p4 Statistics of frequency change between genotypes has been calculated from n=3 mice of each genotype (see Figure 2: supplementary figure 2C and Supplementary Table 3). (**C**) HFSC localization (Sox9, red) in the unwounded (UW) and wound proximal region of WT and C1” “mice at 16 and 24 hours post wounding. Boxed region marks the infundibulum. “*” indicates the wound edge. (**D**) Probability distribution of HFSCs along hair follicles at 16 and 24 hours post wounding in WT and C1^-/-^ wounded and unwounded (UW) skin (see Figure 2: Supplementary Figure 2D and Supplementary table 1). (**E**) Quantification of migrated HFSCs into the epithelial lip at 24 hours post wounding in WT (black, n = 5) and C1^-/-^ (purple, n = 4) wounds. (**F**) Expression analysis of HFSC specific migration genes at 8 hours post wounding in WT (red, n = 6) and C1^-/-^ (purple, n = 4) mice in comparison to C8cKO (pink, n = 5) and C8cKO/C1^-/-^ skin (maroon, n = 3). For PCR each biological replicate was repeated with three technical replicates. Error bars represent S.E.M. “n” denotes biological replicates. “*” indicates p<0.05, “**” indicates p<0.01, “n.s.” indicates “not significant”.

We further validated whether these observations from the genetic wound-healing model, recapitulate a similar phenomenon in excisional wounds in the C1^-/-^ mice. In mice lacking caspase-1, wound-healing is slower and epidermal thickness is inhibited (Lee et al., 2015). Similar to wild-type mice, the largest proportion of HFSCs in the C1^-/-^ mice are present in the bulge region (150-250 μm from the epidermis) under homeostatic (unwounded) conditions (Figures 2C). Whereas the wild-type skin exhibits significant migration of HFSCs into the infundibulum (< 50 μm from the epidermis) within 16 – 24 hours post-wounding, we do not observe significant migration of HFSCs in the skin of wounded C1^-/-^ mice (Figure 2D, Figure 2-figure supplement 1D). The defect in migration is even more pronounced in the C1^-/-^ background at later timepoints as the HFSCs begin to populate the epidermis. At 24 hours post wounding, in the wild-type skin, the number of HFSCs in the epidermis proximal to the wound edge is 2.5 times higher than distal to the wound edge (500 μm from the wound bed) (Figure 2E) but such an efficient migration of HFSCs into the epidermis is not observed in the C1^-/-^ wound. This is consistent with many reports that epithelial stem cell activity is spatially restricted to an area immediately adjacent to the site of tissue damage (Aragona et al., 2017; Ito et al., 2005; Mascré et al., 2012). A similar phenomenon was observed using a lineage tracing approach marking Lgr5 expressing HFSCs, which occupies the lower bulge compartment (Jaks V et al., 2008). The Lgr5 labeled HFSCs exhibited significant migration of HFSC in wounded WT skin compared to wounded skin in the caspase-1 null background (Figure 2-figure supplement 3). Altogether this data demonstrates that the robust migration of bulge HFSCs into the epidermis during wound-healing is critically dependent upon the presence of caspase-1. We next examined whether the effect of caspase-1 on HFSC migration is a manifestation of its impact on the same migratory genes that are upregulated in both the genetic and excisional wound healing models (Figure 1G). Expression of several of the migration associated genes is reduced in the C8cKO/C1^-/-^ epidermis and the C1^-/-^ wound when compared to the C8cKO epidermis and WT wound respectively (Figure 2F). Interestingly, the upregulation of these migration associated genes recover at later timepoints, thereby demonstrating a subsequent, caspase-1 independent phase of HFSC migration (Figure 2-figure supplement 4A). Consistent with this, though there is a significant delay in early stages of HFSC emigration to the epidermis we ultimately observe a large number of epidermal HFSCs by 3 days post wounding in the C1^-/-^ mice (Figure 2-figure supplement 4B and C). This suggests that caspase-1 is a critical initial catalyst of HFSC homing into the epidermis that is further supplemented by alternative signaling pathways to reinforce stem cell chemotaxis into this tissue.

### HFSC migration to the epidermis is mediated by soluble chemotactic cue

An important question that arises from this observation is the mechanism by which Caspase-1 regulates HFSC migration. The canonical function of Caspase-1 as an inflammatory molecule is well studied in immune cells as well as in non-myeloid cells such as epidermal keratinocytes (Sun and Scott, 2016; Yazdi et al., 2010). Caspase-1 is expressed as a zymogen which undergoes activation upon binding to cytosolic complexes known as inflammasomes. The activation of Caspase-1 is required for the cleavage and release of cytokines and several molecules that mediate diverse cellular processes such as survival, inflammation and tissue repair (Keller et al., 2008). We investigated whether it is this Caspase-1 dependent secretome from stressed keratinocytes that is regulating the HFSC migration observed in both the excisional and genetic model of wound-healing. To test if soluble chemotactic factors released from wounded-keratinocytes regulate HFSC migration, we utilized a transwell migration assay with wild-type HFSCs (Wu et al., 2011) and scored their chemotactic index in response to conditioned media (CM) prepared from either wild-type or C8cKO epidermis. In this regard, the genetic model of wound healing is advantageous given that nearly all hair follicles are exhibiting HFSC chemotactic behavior, whereas the excisional wound model only exhibits this phenomenon in a restricted number of follicles. As a consequence, the likelihood of identifying a secreted HFSC chemotactic factor that has heretofore stymied the field is much higher. In comparison to wild-type epidermal CM, C8cKO CM could significantly enhance HFSC migration (Figure 3A). However, epidermal CM from C8cKO/C1^-/-^ mice did not exhibit any HFSC migration promoting activity. This result supports the hypothesis that soluble cues released from wounded keratinocytes stimulates HFSC migration in a Caspase-1 dependent manner. Importantly, the chemotactic cues found in the C8cKO CM does not appear to be a generic stimulus for epithelial stem cell migration as the CM does not enhance chemotaxis of primary mouse epidermal keratinocytes (Figure 3-figure supplement 1B). This suggests that the factor(s) controlling epidermal homing of HFSCs are unique from the classical regulators of epidermal keratinocyte motility (Seeger and Paller., 2015). This unconventional mechanism partly explains the elusive nature of understanding wound-induced HFSC chemotaxis.

**Figure 3:**
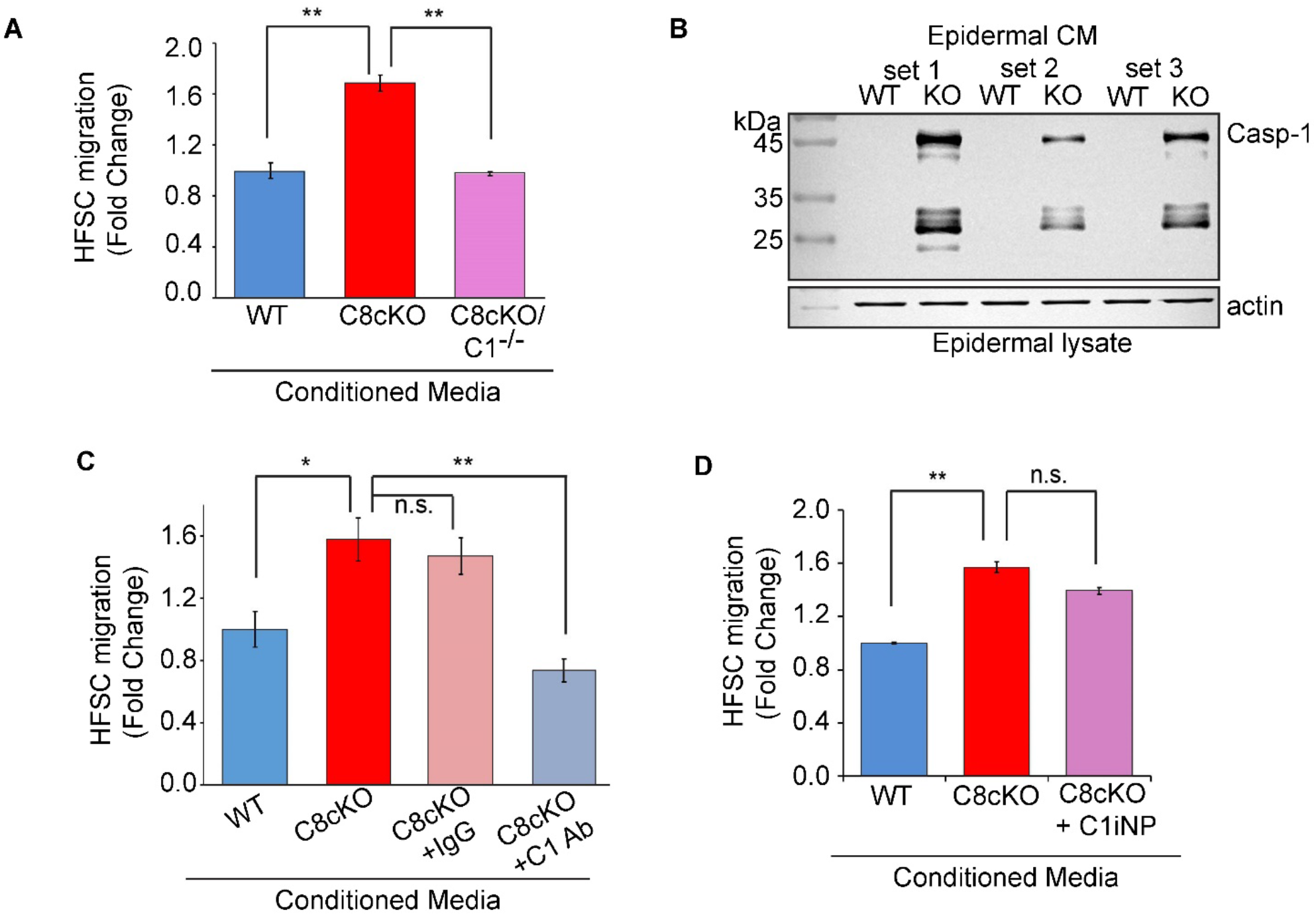
HFSC migration to the epidermis is mediated by soluble chemotactic cues. **(A)** HFSC migration in a Boyden chamber/transwell assay with conditioned media from C8cKO epidermis (C8cKO, red), C8cKO-Caspase-1 null epidermis (C8cKO/C1^-/-^, orange) in comparison to that from wild-type epidermis (WT, gray). Data representive of 4 biological replicates. (**B**) Caspase-1 is detected in conditioned media prepared from C8cKO but not from WT epidermis. The epidermal lysate is probed for actin to ensure equal loading. Three sets of each genotype are shown. **(C)** HFSC migration in response to epidermal conditioned media from wild type (WT, gray) and C8cKO (red) epidermis. Conditioned media from the C8cKO epidermis was immunodepleted with IgG control antibody (C8cKO+lgG, orange) or caspase-1 antibody (C8cKO+C1 Ab, yellow). Data is the mean of 4 biological replicates. (**D**) HFSC migration in response to inhibition of catalytic activity of extracellular caspase-1 present in the C8cKO epidermal conditioned media (C8cKO +C1iNP, pink) compared to vehicle treated C8cKO (red) and WT (gray) epidermal conditioned media. Data is the mean of 3 biological replicates. Error bars represent S.E.M. “*” indicates p<0.05, “**” indicates p<0.01, “n.s.” indicates “not significant”.

Keratinocyte stress leads to Caspase-1 activation and release of IL-1α, which we previously reported to be important to stimulate both epidermal stem cell and HFSC proliferation (Lee et al., 2009, Lee et al., 2017). IL-1α mediates the reciprocal interactions of epidermal keratinocytes and dermal fibroblasts to induce proliferation of keratinocytes of the basal layer of the epidermis (Lee et al., 2009). We have also found that IL-1α mediates the crosstalk between epidermal keratinocytes and resident dendritic epidermal T-cells that ultimately results in the stimulation of HFSC proliferation (Lee et al., 2017). Despite the importance of IL-1α signaling on the proliferation of different epithelial stem cell pools in the skin, we previously observed that it does not affect the migration of the HFSCs into the epidermis (Lee et al., 2017). In light of the fact that the previous tests of the usual candidates of chemotactic factors did not impact directed cell migration of HFSCs, we focused on the often overlooked fact that Caspase-1 activation also results in its own unconventional secretion (Baroja-Mazo et al., 2014; Keller et al., 2008; Lee et al., 2009; Shamaa et al., 2015). This can be in response to stress stimuli such as UV irradiation or wounding of keratinocytes (Feldmeyer et al., 2007; Keller et al., 2008; Lee et al., 2009). Though the secretion of Caspase-1 is a well-documented phenomenon, the function of this extracellular Caspase-1 is largely unknown. We found that epidermal keratinocytes from the C8cKO skin are a major source of extracellular Caspase-1 which can be measured in the conditioned media (Figure 3B). Upon wounding of mouse skin, we observed that extracellular Caspase-1 is found near the wound (Figure 3 – figure supplement 1C) or when we mimicked the wound-reaction in human keratinocytes by simply knocking down caspase-8, the cells responded by increasing the release of Caspase-1 into the culture media (Figure 3 – figure supplement 1D). We probed whether the presence of extracellular Caspase-1 indicates an unknown activity of the molecule in mediating the extensive network of intercellular crosstalk that would induce HFSC chemotaxis in a wound microenvironment. To test this hypothesis, C8cKO-CM was immunodepleted of Caspase-1 and its effect on HFSC migration was investigated (Figure 3C). Resonating with the knockout data (Figure 3A), we find that the removal of extracellular Caspase-1 from the CM abrogates its ability to induce HFSC migration (Figure 3C). The intracellular role of Caspase-1 as part of the inflammasome to mediate unconventional secretion of proteins is primarily dependent on its catalytic function (Keller et al., 2008). Hence, we tested whether the catalytic activity of extracellular Caspase-1 present in C8cKO-CM is required for HFSC chemotaxis. Upon inhibiting Caspase-1 activity in the conditioned media using a cell impermeable pharmacological inhibitor, we found no reduction in migration (Figure 3D). Altogether, these results suggest that extracellular Caspase-1 has a novel, catalytic independent role in promoting the chemotactic migration of HFSCs.

### Recombinant Caspase-1 is sufficient to induce HFSC migration to the epidermis

Our data demonstrates that extracellular Caspase-1 is necessary to promote the directed migration of HFSCs, but is it sufficient to trigger this process? To test this, we bacterially expressed and purified wild-type murine procaspase-1 (rCasp-1 WT) or mutant Caspase-1 (C284A) to abolish its catalytic activity (rCasp-1 C284A) (Broz et al., 2010) (Figure 4 – figure supplement 1A). The identity of the recombinant proteins was confirmed by Western blotting for Caspase-1 and their catalytic activity was verified by a substrate cleavage assay (Figure 4 – figure supplement 1B). Whereas the purified rCasp-1 WT was efficient in cleaving its substrate, the rCasp-1 C284A is indeed catalytically dead. We then tested whether the wild-type or the mutant protein is sufficient to stimulate HFSC chemotaxis. In comparison to control treated samples, both the rCasp-1 WT and rCasp-1C284A proteins had equivalent capability to induce chemotaxis of HFSCs (Figure 4A). This supports our earlier observation based on pharmacological data that the catalytic function is dispensable for Caspase-1 to operate as a HFSC chemoattractant (Figure 3D). Moreover, the ability of Caspase-1 to induce migration in HFSCs is independent of cellular proliferation, as measured by the response of mitomycin C treated HFSCs to rCasp-1 C284A in comparison to untreated HFSCs (Figure 4B).

**Figure 4:**
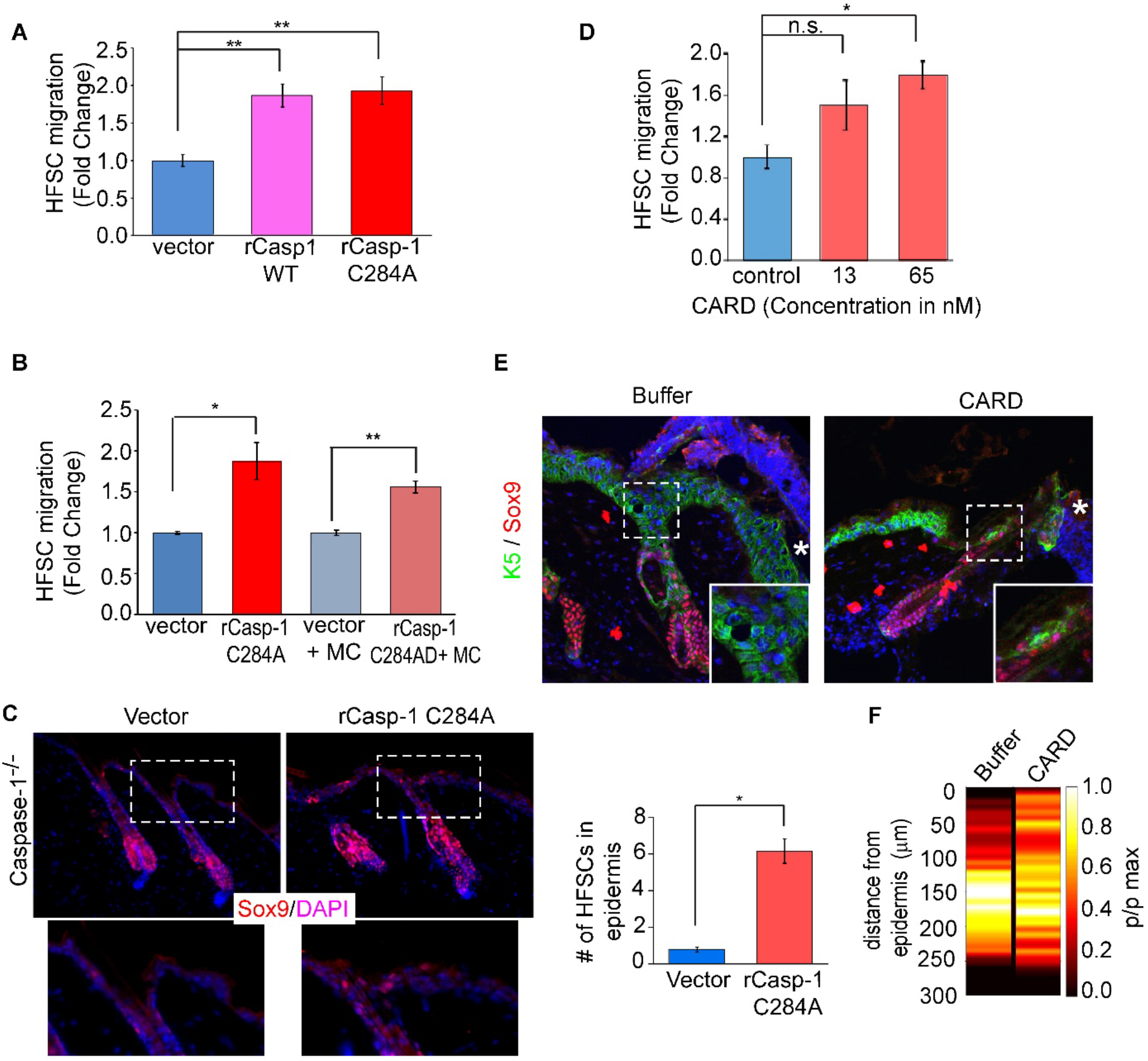
Recombinant Caspase-1 is sufficient to induce HFSC migration into the epidermis. **(A)** HFSC chemotaxis to recombinant Caspase-1 proteins; wild-type (rCasp-1 WT, pink), catalytic dead (rCasp-1 C284A, red) in comparison to vector proteins control (vector, blue). n= 4. (**B**) Chemotaxis of proliferating and nonproliferating (upon Mitomycin treatment, +MC) HFSCs to rCasp-1 C284A in comparison to vector control, n=3. **(C)** Immunofluorescence images of HFSC localization (Sox9, red) in caspase-1 null skin explant treated with rCasp-1 C284A or vector control. Top panel is a representative image, with magnified insets, and the bottom panel is the quantification of number of HFSCs in the epidermis within a 100 μm^2^ area. Quantification is based on 3 biological replicates. **(D)** Analysis of HFSC chemotaxis to recombinant Caspase-1 CARD (red) in comparison to buffer control (blue) from three biological replicates. Data shows mean and S.E.M. for all experiments. (**E**) Representative immunofluorescence images showing the migration of HFSCs to the infundibulum upon application of recombinant CARD or buffer within 24 hours of wounding on caspase-1 null mouse skin. Insets are magnified views of the dashed boxes. “*” marks the wound edge. **(F)** Probability distribution of HFSCs in wound-proximal hair follicle from buffer or CARD treated wounds (see Figure 2: Supplementary Figure 4F and Supplementary table 5). “n” denotes biological replicates. “*” indicates p<0.05, “**” indicates p<0.01, “n.s.” indicates “not significant”.

This complements the observations made in Figures 1 and 2, where we found that the early migratory wave (at 16 - 24 hours) of HFSCs in a Caspase-1 dependent manner is independent of proliferation in the niche. Moreover, this finding is consistent with recent reports that during the re-epithelialization process of wound healing, migrating cells and proliferating cells are two spatially distinct populations (Aragona et al., 2017; Park et al., 2017). Extending on these *in vitro* observations, we assayed for the capacity of rCasp-1 C284A to induce HFSC migration to the epidermis in skin explants from C1^-/-^ mice. Indeed, we find that there is relocalization of HFSCs to the interfollicular epidermis upon treatment with rCasp-1C284A (Figure 4C). Given this unusual catalytic-independent function of Caspase-1 as a chemoattractant, we next investigated which domain is responsible for this novel activity. Caspase-1 is comprised of an N-terminal Caspase Activation Recruitment Domain (CARD), followed by the p-20 and p-10 domains (Broz et al., 2010). Autoproteolysis results in the separation of these individual domains, followed by the heterodimerization of the p20 and p10 domains into the catalytically active form. Since the catalytic activity is not required for inducing HFSC migration (Figure 3D and Figure 4A), we focused on the CARD which is the protein-protein interaction region of Caspase-1 (Park, 2019). The murine Caspase-1 CARD was tagged with polyhistidine, bacterially expressed and purified by affinity chromatography (Figure 4 – figure supplement 1C) and its folding status was confirmed by circular dichroism measurement whereby its spectra (Figure 4 – figure supplement 1D) matches largely to the published spectra of CARDs from other proteins (Palacios-Rodríguez et al., 2011). We also observed that the recombinant CARD was able to bind to the bulge region of the hair follicle where the HFSCs reside (Figure 4 – figure supplement 1E). Chemotaxis assays with recombinant Caspase-1 CARD showed enhanced migration of HFSCs (Figure 4D) but did not have an effect on primary epidermal keratinocytes (Figure 4 – figure supplement 1H). Moreover, the intracellular pool of Caspase-1 null HFSCs exhibited enhanced migration in the transwell migration assay in response to recombinant CARD indicating that the cytosolic Caspase-1 is not required for migration (Figure 4 – figure supplement 1G). Finally, we examined whether exogenous application of CARD to the excisional wounds on caspase 1 null mice can enhance HFSC migration. Within 24 hours of wounding, CARD application increased the mobilization of Sox9+ HFSCs into the infundibulum of wound proximal hair follicles (Figure 4E and 4F). Consistent with this, Lgr5 labelled HFSCs demonstrated that CARD application on the wounds of caspase-1 null mice significantly enhanced cell migration (Figure 4 – figure supplement 2). These results indicate that the extracellular Caspase-1 mediates its non-canonical chemotactic function largely by its CARD.

### Caspase-1 is a regulator of stem cell responses in the skin during UV damage

Our data demonstrates that wounding leads to the release of extracellular Caspase-1 from stressed keratinocytes, which acts as a chemotactic factor to elicit HFSC migration to the epidermis. However, this is not the only scenario reported to exhibit the secretion of Caspase-1. Skin carcinomas (Drexler et al., 2012; Karki and Kanneganti, 2019) and other inflammatory skin diseases such as psoriasis (Johansen et al., 2007), atopic dermatitis (Antonopoulos et al., 2001; Li et al., 2010) and sunburn (Faustin and Reed, 2007), are all marked by Caspase-1 activation.

Thus we examined whether extracellular Caspase-1 mediated HFSC migration into the epidermis is a conserved phenomenon that may explain the epithelial hyperplasia associated with these pathologies. In particular, we investigated whether UV irradiation could induce HFSC migration *in vitro* and *ex vivo*. Since UV irradiated mouse keratinocytes release Caspase-1 (Feldmeyer et al., 2007; Keller et al., 2008), we tested whether conditioned media prepared from UV irradiated epidermal keratinocytes is capable of stimulating HFSC chemotaxis. We found that CM of UV irradiated keratinocytes can significantly enhance HFSC chemotaxis compared to CM from non-irradiated keratinocytes (Figure 5A). We also examined whether UV irradiation of mouse skin explants displayed epidermal homing behavior of HFSCs in a Caspase-1 dependent manner. Within 16 hours of UV irradiation of mouse skin explants, we observed HFSCs migrating upwards along hair-follicles to the epidermis in wild-type skin (Figure 5B). However, there was a substantial reduction in the number of HFSCs homing into the epidermis in C1^-/-^ skin explants exposed to UV irradiation (Figures 5B and 5C).

**Figure 5:**
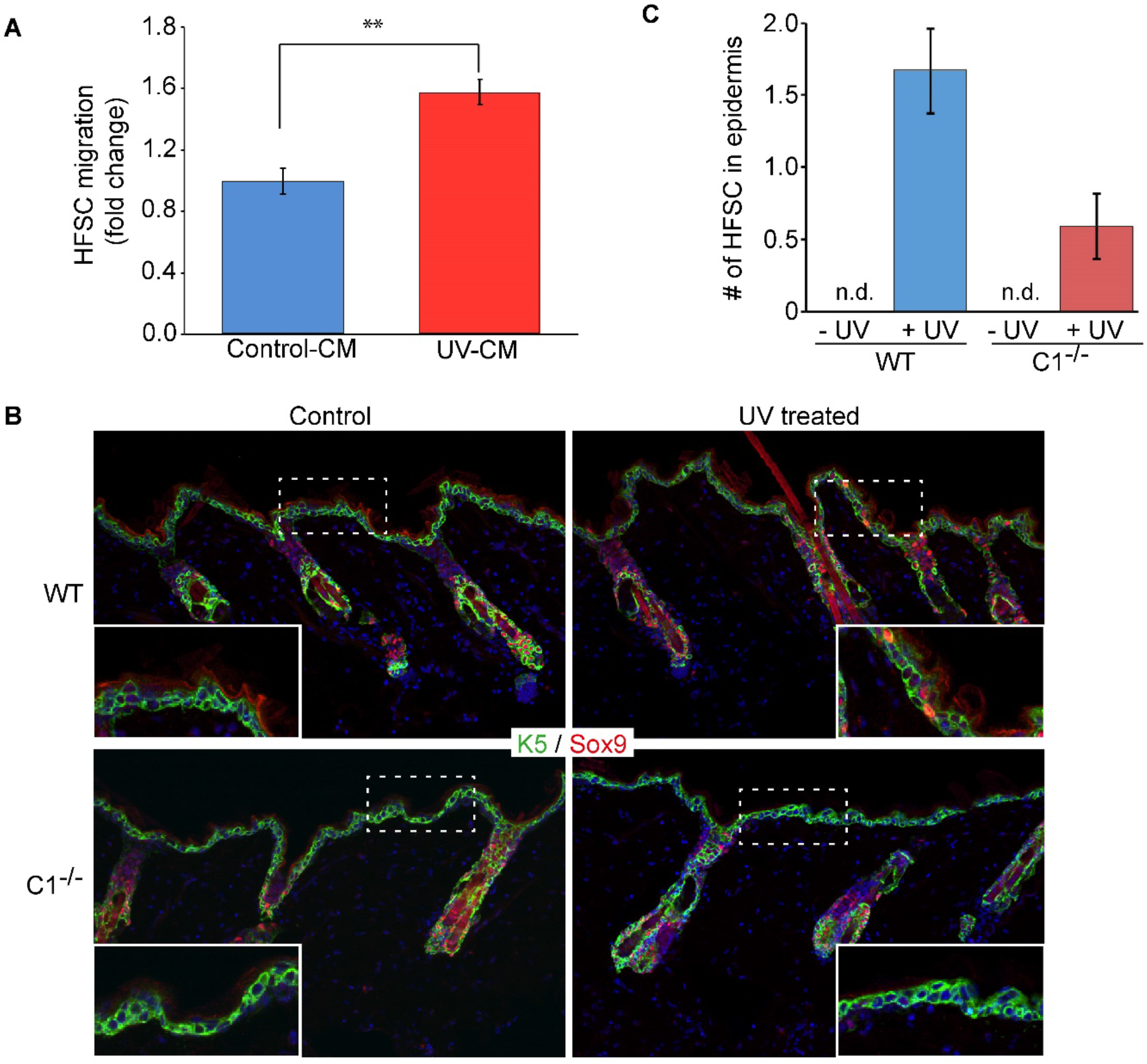
Caspase-1 mediates UV irradiation induced HFSC migration to the epidermis. **(A)** HFSC migration in response to conditioned media from control (Control-CM, blue) or UV treated (UV-CM, red) primary mouse keratinocytes. n=3. (**B**) Immunofluorescence images of HFSC migration towards the epidermis visualized in WT and C1^-/-^ skin explants in untreated (control) or UV treated conditions with Sox9 staining (red). Inset shows magnified view of boxed region of interfollicular epidermis. **(C)** Quantification of HFSCs in the epidermis in the control and UV treated skin explants from WT (gray) and C1^-/-^ (purple) mice, n=3 of each genotype, n d = not detected. Data shows mean and S.E.M. of all experiments, “n” represents biological replicates. “**” indicates p<0.01

## DISCUSSION

Altogether a model emerges wherein Caspase-1 is an important mediator of both inflammatory signaling as well as stem cell migration in wound-healing and other skin pathologies characterized by inflammation. This novel chemotactic function is mediated by the extracellular Caspase-1, more specifically by its protein-protein interaction domain CARD. Our work elucidates the role of extracellular Caspase-1 as a bulge specific HFSC chemoattractant but it might have similar roles on other stem cells in skin appendages known to participate in wound-healing.

Mobilization of multiple cutaneous stem cell pools is critical for injured tissue to reestablish the critical barrier function of the skin. Herein, we have probed the mechanism by which a specific class of multipotent epithelial stem cells resident in the hair follicle niche, are repurposed to form epidermal keratinocytes when there is a wound in the skin. Under homeostatic conditions, these hair follicle stem cells (HFSCs) are activated periodically for hair follicle regeneration, during which they proliferate and their progeny migrate into the dermis to form a new hair follicle (Myung and Ito, 2012; Rompolas and Greco, 2014). Upon wounding, however, the HFSCs are redirected towards the epidermis, whereupon they differentiate to form epidermal keratinocytes (Gonzales and Fuchs, 2017; Plikus et al., 2012). The signals regulating HFSC driven homeostatic hair regeneration have been well studied but the cues driving HFSC wound response have been difficult to identify. This disparity in our understanding of these two behaviors arose as a result of technical challenges. The identification of the wound cues and cellular responses occurring only within the small region surrounding the wound site yielded a low signal to noise ratio. Our ability to tap into a genetic model of wound-healing, the C8cKO mouse, which exhibits a wound-healing response throughout the skin, allowed us an amplified view of the process both at the level of the source of the wound-cue (keratinocytes) and the responding cells (HFSCs).

By validating the insights garnered from the genetic model with excisional wounds, we have identified a bona fide cue that drive the HFSCs to the wounded epidermis (Figure 6). A model emerges wherein Caspase-1 is an important mediator of both inflammatory signaling as well as stem cell migration in wound-healing and other skin pathologies characterized by inflammation. This novel chemotactic function is mediated by the extracellular Caspase-1, and, in particular, its protein-protein interaction domain CARD. These observations build upon our previous findings that the downregulation of epidermal caspase-8 is a normal phenomenon upon damage to the skin (Lee et al., 2009). One consequence of the reduction in caspase-8 expression is the activation of the Caspase-1 containing inflammasome in the epidermal keratinocytes. This launches the secretion of the reservoir of IL-1α from wounded keratinocytes that mediates the cellular cross-talk between the epidermal, dermal and immune cell populations in the skin (Lee et al., 2015). Herein, we found that Caspase-1 itself is released from stressed keratinocytes in multiple scenarios, such as upon excisional wounding, the loss of epidermal caspase-8 in neonates as well as adults, or UV irradiation, into the extracellular space. This is in line with other reports that Caspase-1 is released from both immune cells (Baroja-Mazo et al., 2014; Sarkar et al., 2009) and keratinocytes (Keller et al., 2008; Lee et al., 2009) by the unconventional secretory pathway. Immune cells release Caspase-1 and associated inflammasome complex proteins via exosomes that deposit these activated proteins into the cytoplasm of nearby immune cells to stimulate the activation of their inflammation-mediating machinery (Baroja-Mazo et al., 2014; Sarkar et al., 2009). As such, it is an efficient mechanism of amplifying the number of immune cells generating an inflammatory response. Keratinocytes upon UV stress release extracellular Caspase-1 and a host of other proteins in a manner that depended upon Caspase-1’s catalytic activity (Keller et al., 2008). The ejection of these proteins from the stressed cells is speculated to serve the purpose of cytopreservation and escaping cell death but the role they would play in the extracellular space was not considered. Though the exact mechanism of wound-induced Caspase-1 release from keratinocytes remains undefined, the ability of soluble recombinant protein expressed in bacteria to induce chemotaxis of HFSCs suggest that this protein is not packaged completely within an exosome as reported for some myeloid cells.

**Figure 6:**
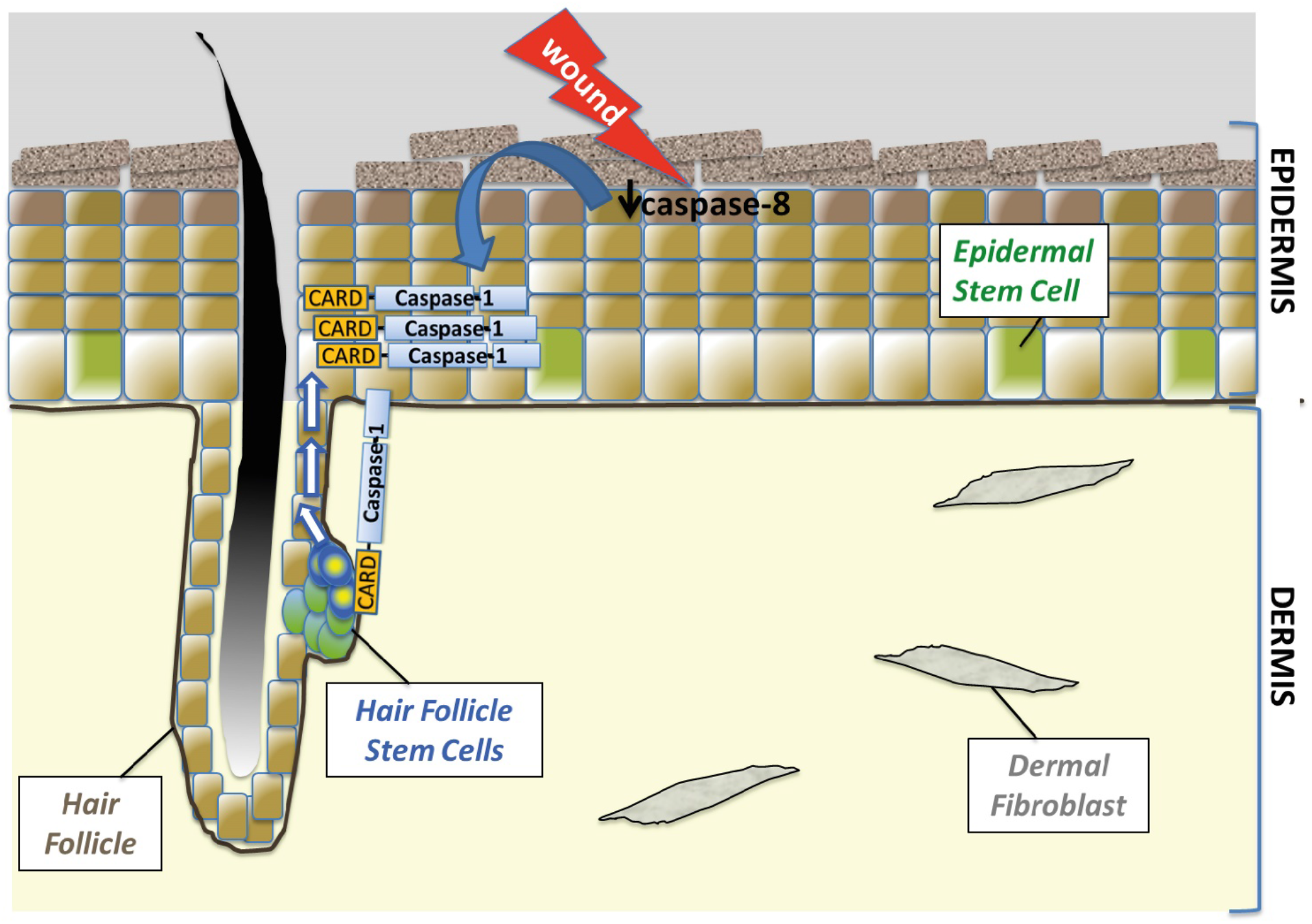
Model of Caspase-1 mediated HFSC homing to the epidermis upon wounding. Injury to the skin leads to the downregulation of caspase-8 in the upper (granular) layer of the epidermis. The reduction of caspase-8 leads to the secretion of caspase-1 from the keratinocytes. Via caspase activation and recruitment domain (CARD) of caspase-1, hair follicle stem cells are activated to migrate to the wounded epidermis.

Interestingly, we have previously shown that there is no evidence of cell death in the caspase-8 conditional knockout at the ages analyzed in the manuscript (Lee et al., 2009; Li et al., 2010). This is consistent with our observation that the barrier function of the epidermis (measured via transepidermal water loss and dye penetration assay) is not compromised in the neonatal mouse. There are lesions that form in the caspase-8 conditional knockout mouse but these occurred after 3 months (Li et al., 2010). As such, the presence of Caspase-1 in the extracellular milieu is an active process and not simply leaking out of dying cells as has been previously reported (Feldmeyer et al., 2007; Keller et al., 2008). Instead, our results are consistent with several papers reporting the unconventional secretion of Caspase-1 from activated monocytes and macrophages without cell lysis (Feldmeyer et al., 2007; Keller et al., 2009, 2008).

Caspase-1 has a N-terminal protein-protein interaction domain, CARD, similar to other inflammatory Caspases (Lavrik and Krammer, 2009), which interacts with other CARD containing proteins in the cytosol. In fact, the activation of procaspase-1 to active Caspase-1 is dependent on this CARD-CARD interaction of Caspase-1 to ASC leading to the formation of the inflammasome complex. Caspase-1 activation is marked by the cleavage of procaspase-1 into CARD, p-20 and p-10 domains, the latter two forming the heterodimeric active protease. Interestingly, we find both procaspase-1 as well as cleaved versions of the protein in the extracellular space. Prompted by the observation that chemotactic activity is independent of catalytic function of the Caspase-1, we probed whether the CARD of Caspase-1 could harbor the novel chemotactic function of this protein. Indeed, we found that recombinant Caspase-1 CARD could elicit HFSC migration but not that of primary keratinocytes. We have demonstrated the responsiveness of Sox9 positive HFSCs and an endogenous marker Lgr5 located in the bulge to CARD, but whether other resident hair follicle stem cells identified by endogenous markers such lgr6 (upper bulge HFSCs), and lrig1 (infundibulum HFSCs) are receptive to this signal is yet to be determined.

Beyond the current discovery of the chemotactic cue for HFSC migration, the downstream signaling cascade(s) leading to directional migration remains open for investigation. An obvious question is the cognate receptor for the CARD containing ligand. A cell surface interaction to initiate the migratory behavior is postulated given the fact that we have not been able to observe the intracellular uptake of His-tagged recombinant CARD in the recipient HFSC (data not shown). In general, a downstream effect of this directional cue is the polarization of the HFSC cytoskeleton and its associated migratory machinery (Ridley et al., 2003; Seetharaman and Etienne-Manneville, 2020). In epithelia such as the epidermis and hair follicle, cell-cell junctional and ECM interactions must be dynamically remodeled in coordination with the polarizing machinery leading to efficient migration towards the wound-bed. There is evidence for the role of GSK-3β in regulating the polarization of microtubule-actin interactions through the binding/unbinding of ACF7, a cytoskeletal crosslinker in HFSCs, that affect their entry into the wound (Wu et al., 2011). It remains to be tested whether Caspase-1 CARD mediated chemotactic migration utilizes this same mechanism.

It is noteworthy that even though the loss of caspase-1 significantly delays the epidermal homing of HFSCs, it nevertheless does not completely abrogate this phenomenon. This suggests that extracellular Caspase-1 might be an early chemotactic cue while other regulators of chemotaxis potentiate this effect, or can compensate in instances where the protein is absent. Given the importance of wound healing for the health of the organism, it is not surprising that many of the processes in the repair program have redundant mechanisms to ensure the completion of this critical process. It is important to note that even though the caspase-1 null animal used in our studies (Li et al., 1995) is also deficient in caspase-11 (Kayagaki et al., 2011), recombinant Caspase-1 or CARD is sufficient to restore hair follicle stem cell migration in the caspase-1 deficient mouse. Moreover, caspase 4 and caspase-11 are not in the conditioned media generated from the C8cKO, which has promigratory activity (data not shown).

This mechanism of homing of HFSCs to the epidermis is not only important in physiological scenarios but offers important insights into pathological conditions. For instance, tumorigenesis and cancer metastasis usurp natural cellular responses observed during wound-healing such as proliferation and migration in a deregulated manner. Indeed, chronic wounds and chronic inflammatory stress such as sunburn and UV damage can push normal stem cells towards malignant transformation. In the skin, carcinomas may arise from both epidermal and hair follicle stem cells typically due to oncogenic mutations in the stem cell compartment but upon prolonged inflammation or injury (such as that associated with repeated UV damage or sunburn) the stem cells are transformed to cause carcinogenesis (Singh et al., 2013; White et al., 2011; White and Lowry, 2014). In squamous cell carcinoma and basal cell carcinoma of the skin, HFSCs hyperproliferate and migrate to the epidermal compartment, but extrinsic drivers of this deregulated behavior is unknown. Our observation that UV damage to the skin can cause HFSC migration to the epidermis in a caspase-1 dependent manner, suggests that this chemotactic cue driving the wound-response of HFSCs could be common to pathological transformation in the skin observed upon sunburn and other inflammatory and hyperproliferative skin pathologies such as atopic dermatitis and psoriasis.

## MATERIALS AND METHODS

### Mouse models

Caspase-8 conditional knock-out is created by K14-Cre driven knock-out of caspase-8 in the epidermis (Lee et al., 2009, 2017). The caspase-1 null mice have been described before (Lee et al., 2015; Li et al., 1995). The caspase-8 inducible knock-out is K14-CreER driven in B6/J background (Vasioukhin et al., 1999). K14-CreER, C8 ^fl/fl^/C1^-/-^ mice were created by crossing caspase-8 inducible knock-out is K14-CreER driven in B6/J background and caspase-1 null mice. Adult mice (7 week old) mice were injected with 6 mg of Tamoxifen in Corn Oil (Sigma) intraperitoneally for 3 days, with gaps of alternate days. Following a wait period of 2 weeks, the mice were sacrificed for histological analysis of the skin. Lgr5 eGFP tdT mice were created by crossing B6.129P2-Lgr5^tm1(cre/ERT2)Cle^/J (JAX#008875) with B6.Cg-Gt(ROSA)26Sor^tm14(CAG-tdTomato)Hze^/J (JAX# 007914). Lgr5 eGFP tdT caspase1-/- mice were created by mating Lgr5 eGFP tdT mice with caspase1 null mice. Adult mice were injected with 1.25mg of Tamoxifen in Corn Oil (Sigma) intraperitoneally for 5 consecutive days. Following a waiting period of one week, the mice were used for the wounding experiments and later sacrificed for histological analysis of the skin. Animal work conducted was approved by the inStem Institutional Animal Ethics Committee following norms specified by the Committee for the Purpose of Control and Supervision of Experiments on Animals, Government of India.

### Wounding and explant treatment experiments

8 week old mice were shaved and wounded under isoflurane induced anesthesia using 5 mm punch biopsy to make full thickness excisional wounds. Animals were sacrificed at specific timepoints to collect skin samples for histological analysis. For obtaining skin explants, caspase-1 null mice were sacrificed and 1 cm^2^ piece of skin was excised. After removal of subcutaneous fat by scraping, the explant was floated on DMEM-10% FBS media for the duration of the experiments.

### Cell culture and conditioned media preparation

All experiments were approved by the inStem Institutional Biosafety Committee. Wild-type primary mouse keratinocyte cultures are established by isolating cells from the epidermis of newborn pups and cultured in low-calcium (0.05 mM) mouse keratinocyte media (E-media) as described previously (Lee et al., 2009). For the isolation of keratinocytes and for preparation of conditioned media, epidermis is chemically dissociated from the dermis by dispase treatment (1 hour at 37 °C) from skin of newborn pups aged p0 - p3. For primary keratinocyte isolation, the epidermis is briefly treated with 0.25% Trypsin-EDTA, followed by culturing the isolated cells on Mitomycin C treated J2-3T3 feeder cells in low-calcium (0.05 mM Calcium) E-media (Nowak and Fuchs, 2009). For preparation of epidermal conditioned media, the isolated epidermis was floated in serum-free E-media for 16 hours. HFSCs were isolated from 8 week old wild-type mice skin by following standard methods and cultured on feeders for 8 passages in HFSC culture media (E-media with 0.3 mM calcium) (Nowak and Fuchs, 2009) or the 3C culture method reported by Chacón-Martínez CA et al., 2017.

### UV irradiation experiments

Differentiated keratinocyte culture was exposed to 50mJ of UV radiation in phosphate buffer saline following which serum free keratinocyte culture media was added. Conditioned media was collected after 4 hours. The skin explants from WT and C1^-/-^ mice were UV irradiated at 250mJ/cm^2^, following which these were floated in DMEM-10% FBS media for 16 hours before collection and processing.

### Image acquisition and analysis

Fluorescent images of tissue samples were obtained on the Olympus FV3000 confocal microscope at the Central Imaging and Flow Facility at NCBS. For image processing ImageJ software (Fiji) was used. To analyze the distribution of HFSCs along hair follicles, the positions of Sox9+ nuclei and the hair-follicle-epidermis junction was marked manually in Fiji (Figure 1 – figure supplement 2A). In the wounded samples, the first hair follicle immediately adjacent to the wound, particularly within 300µm were considered. The spatial coordinates of all the marked nuclei was exported to MATLAB for further analysis. The MATLAB routine we developed calculates the distance of individual Sox9+ nuclei in a hair follicle from the epidermal junction, then based on data similarly collected from several hair follicles across multiple biological samples, the probability of occurrence of these nuclei along the length of hair follicles is estimated (Figure 1 – figure supplement 2B). Typically, the spatial frequency was calculated from 8 μm bins along the length of hair follicles. For the representation by heatmap, the frequencies were normalized to their maxima and plotted such that white color represents 1 and the darker shades till black represents lower values till 0. Differences in localization frequency were considered significant for p<0.05 with n = 3 mice for each condition, and represented by the shaded box (Figure 1 – figure supplement 2A, E). Details of animals and hair follicle numbers for each set of statistical analysis are provided in Supplementary Table 1, 2, 3, 4, 5 and 6. The MATLAB code used for data analysis of frequency distribution of HFSC migration in the epidermis can be found at the following link: https://github.com/skinlab-sunnyk/HFSCs.

### Cloning, expression and purification of recombinant proteins

The murine Caspase-1 wild-type and catalytic dead constructs were gifted to us in a retroviral vector, pMScV (Broz et al., 2010). The genes for procaspase-1 were cloned into pET-28a vector by engineering flanking Xho-1 and Nco-1 sites. For purification from bacterial lysates, the protein sequence was followed by C-terminal 6x-His tag expression from the vector backbone. The constructs were expressed in BL21 pLysS under IPTG induction at 18°C for 16 hours. The cells were lysed by sonication in lysis buffer (50mM Tris-HCl, 300mM NaCl, 10% glycerol, 5mM DTT, pH 8.0 with EDTA free Protease Inhibitor Cocktail, DNase I and lysozyme). Following centrifugation at 16000g, the supernatant was incubated with Ni-NTA beads for 3 hours at 4°C in the presence of 10mM imidazole. After washing with 20mM and 40mM imidazole, the recombinant proteins were eluted at 120mM imidazole at 4°C in lysis buffer without additives. Following dialysis to imidazole free buffer the proteins were snap-frozen for future use. The CARD of the caspase-1 gene synthesized (Geneart, Thermo) after codon optimization with N-terminal Strep-II tag with Enterokinase cleavage site and C-terminal 10x-His tag, then cloned into pET-28a vector using the same restriction sites. Cell pellet was lysed by probe sonication in a denaturing buffer composed of 50mM Tris-HCl, pH8.0 with 6.5 M GdnHCl, 300 mM NaCl, and 50 mM imidazole; following centrifugation at 16000g, the supernatant was incubated with Ni-NTA beads (Thermo). Following on-column refolding using a reducing Urea gradient in Tris-buffer with imidazole, final elution was performed at 500 mM imidazole at 4°C. Following dialysis into imidazole-free buffer, the proteins were snap-frozen for future use. The purity of the proteins was tested by silver staining of the eluates as well as identity was confirmed by Western blotting for Caspase-1. The concentration of the recombinant proteins was ascertained by both Bradford Reagent and quantification from silver stained gels in comparison to BSA gradient.

### Chemotaxis assays and quantitation

HFSCs or mouse primary keratinocytes were trypsinized and plated onto the top chamber of either 24 well or 96 well transwell plates having 8μm pore sizes (Corning). The conditioned media or proteins being tested were diluted in HFSC culture media or low Calcium E-media and added to the bottom chamber. Cell migration was assayed after 12 hours of incubation when the cells were fixed by PFA and stained with Crystal Violet. The top of the transwell was cleaned before counting the cells in the bottom using the Cell Counter plugin on ImageJ-Fiji.

### RNA isolation and qPCR

Wound proximal tissue was obtained by using an 8mm punch biopsy tool to excise the tissue from a 5mm punch wound. A biopsy was taken 3cm away from the wound as unwounded tissue. Epidermal samples were collected immediately after isolation by dispase treatment. These samples were collected in chilled TRIzol reagent and crushed using a tissue homogenizer (at 8000 rpm). Total RNA was extracted from crushed tissue or cell lysate using the TRIzol reagent protocol (TAKARA/Thermo). cDNA was synthesized using Superscript III (Thermo), followed by quantitative PCR using the Power SYBR mix (Life technologies) in the BioRad CFX384 machine. *Actin* or *18S* expression was used as reference for normalization. Primer sequences used are given in Supplementary Table 7.

### RNA Seq Data analysis

For RNA Seq reads we have taken publicly available data at NCBI repository (GEO accession ID: GSE89928) from the study by (Ge et al., 2017). The raw FASTQ files for two biological replicates, each from control and wound bulge stem cells are processed for further analysis. Briefly, the quality of FASTQ files were analyzed by FASTQC. Trimmomatic tool (Bolger et al., 2014) were used to remove any adapter contamination, and high-quality reads were aligned to Mouse reference genome (GRCm38) using TopHat 2. The data were analyzed further using the “Tuxedo” pipeline: TopHat2⍰ Cufflinks ⍰ Cuffmerge ⍰ Cuffdiff (Trapnell et al., 2012). The significantly differentially expressed genes (q value < 0.05) from Cuffdiff output were used further for Gene Ontology (GO) enrichment analysis using DAVID.

### Reagents

Antibodies used for HFSC isolation are anti-CD49f-PE conjugated (BD 561894) and anti-CD34-FITC conjugated (eBioscience, 11-0341-82). Antibodies used for immunofluorescent staining of tissues are anti-Sox 9 (Abcam, rabbit, 1:100), anti-Ki67 (Abcam, rabbit,1:200), and anti-Keratin 5 (raised in lab, chicken, 1:100), anti-mouse Caspase-8 (Enzo, rat, 1:100). Nuclei were stained with Hoechst. Secondary antibodies used were Alexa-488 anti-chicken (1:400) and Alexa-568 anti-rabbit (1:400) from Life technologies. For immunoprecipitation from conditioned media, Casper-1 antibody (AdipoGen) and corresponding mouse IgG (CST) was used. Antibodies used for Western blot: Caspase-1 (Abcam, rabbit, 1:500), Caspase-8 (CST, mouse, 1:1000), GAPDH (Sigma, rabbit, 1:2000), Caspase-1 (Santa Cruz, mouse, 1:500). Caspase-1 non-permeable (Cat # 400010) inhibitor was obtained from Millipore and used at a concentration of 10μM. Caspase-1 substrate used is Ac-YVAD-AMC (Enzo Lifesciences) at 15μM.

## ACKNOWLEDGEMENTS

We thank Petr Broz (University de Lausanne) for the gift of the caspase-1 constructs. Animal work in the NCBS/inStem Animal Care and Resource Center was partially supported by the National Mouse Research Resource (NaMoR) BT/PR5981/MED/31/181/2012;2013-2016 & 102/IFD/SAN/5003/2017-2018 from DBT. We thank Central Imaging and Flow Cytometry Facility (CIFF) of the Bangalore Life Sciences Cluster for image acquisition support.

## SUPPLEMENTARY INFORMATION

**Figure 1: Figure Supplement 1.**
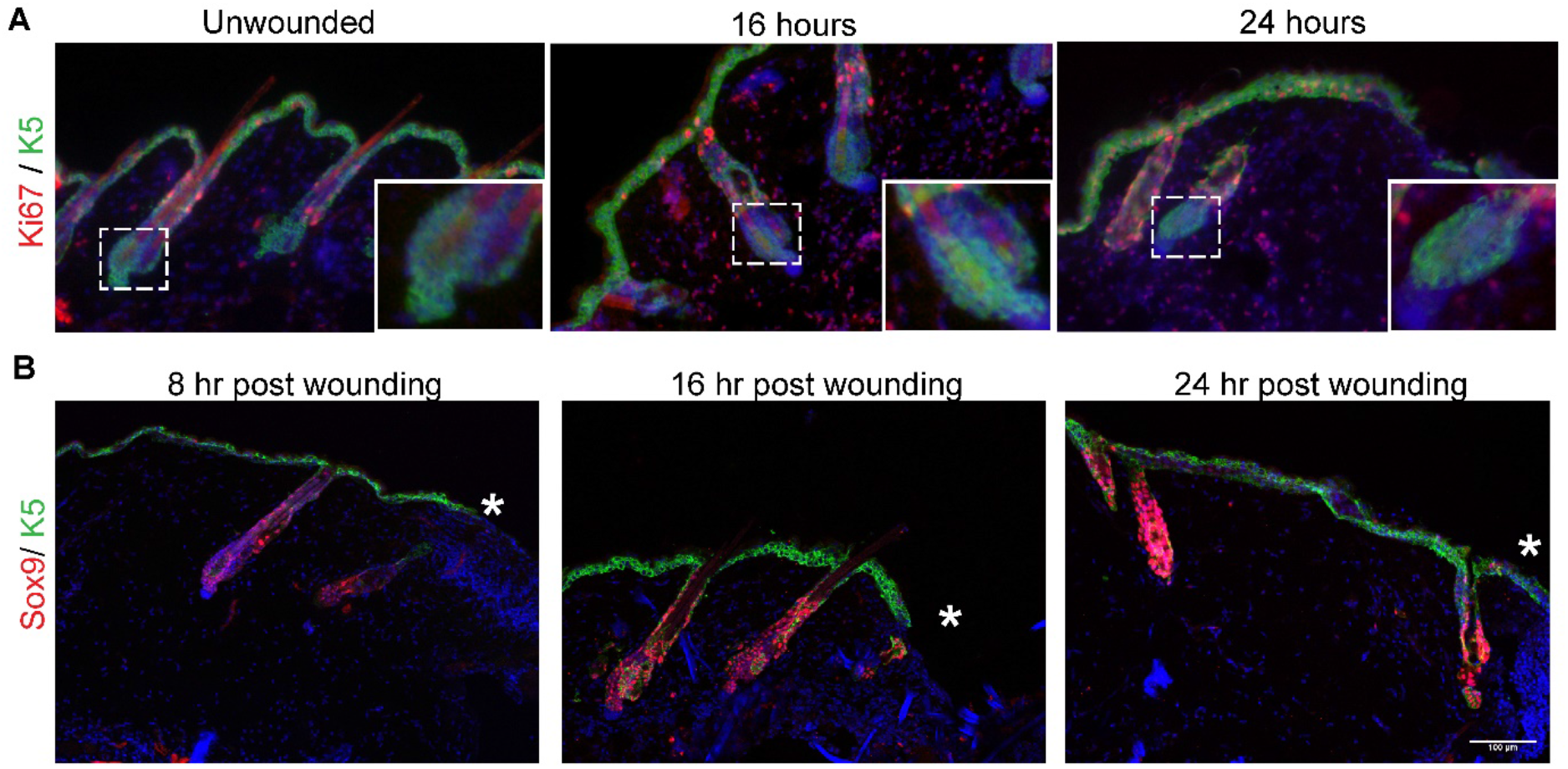
**(A)** Proliferation marked by Ki67 (red) in the unwounded (UW) skin and at 16 and 24 hours of wound-healing (W). Epidermis and hair follicles are marked by K5 (green), and the bulge is shown in the inset. **(B)** Wound proximal skin (* marking the wound-bed) showing typical position of hair follicles analysed for statistics in Figure 1 B, E, at each of the 8,16 and 24 hour time points of wound-healing

**Figure 1: Figure Supplement 2.**
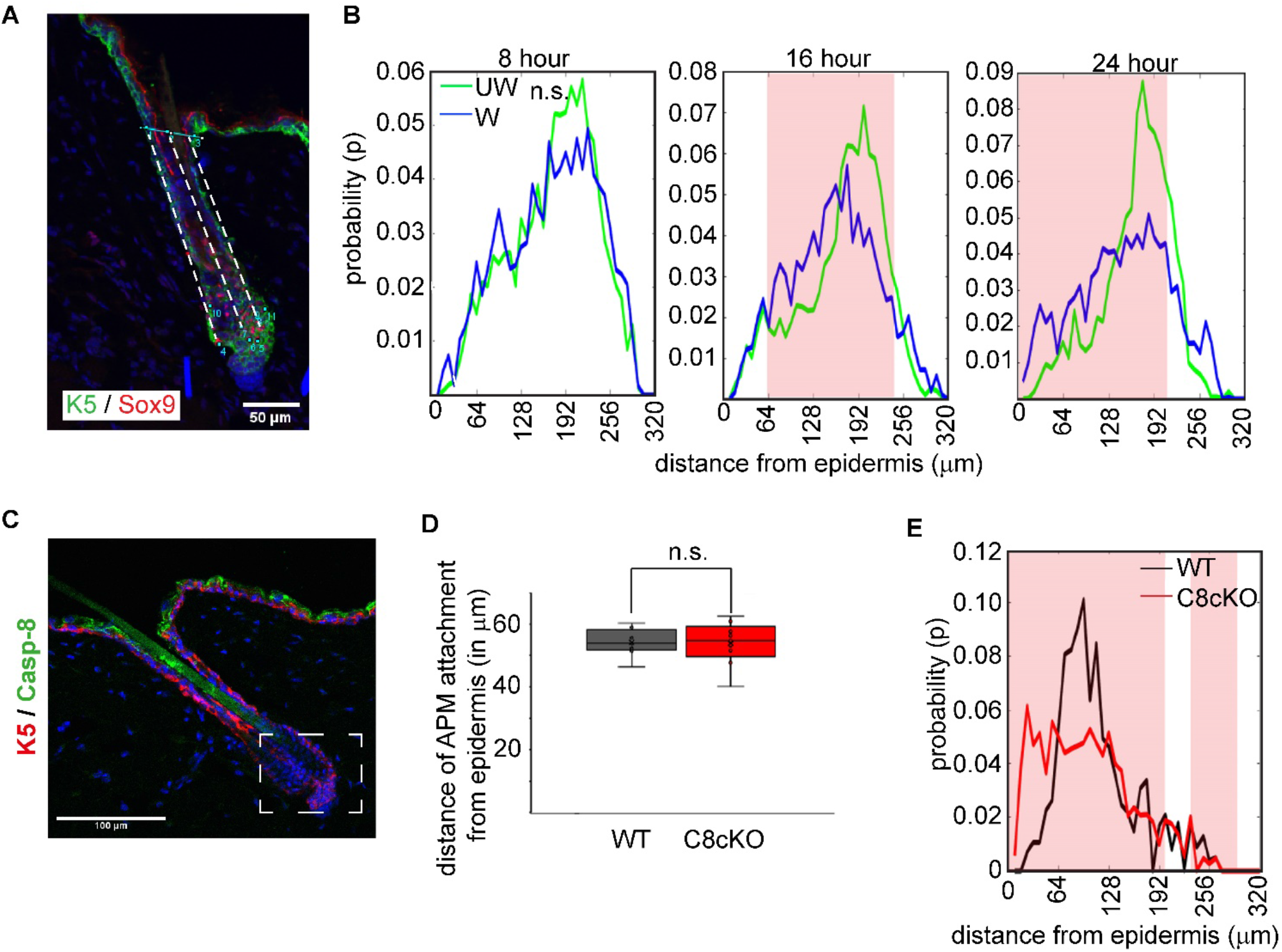
**(A)** HFSCs (Sox9, red) were marked along hair follicles and the epidermal junclion of the hair follicle was marked by 3 reference points. Distance of individual HFSCs were measured from the epidermal junction (dotted lines) and the frequency of localization along the length of the hair follicle was measured (See Methods). Scale bar = 50um. **(B)** Probability plot of HFSC localization measured along hair follicles in unwounded (UW, green line) and wounded (W, blue line) skin at 8, 16 and 24 hours post wounding, wherein the x-axis represents the distance of HFSCs from the epidermis. Pink shaded box marks the region where the probability is significantly different (p <0.05) between UW and W conditions (see supplementary table 1 for sample size of each set). **(C)** Caspase-8 (green) expression in the epidermal compartment is observed in the granular layer and not in the HFSC niche (boxed region). **(D)** In postnatal day 4 (p4) pup skin, the position of the HFSCs is deter mined by the distance of the APM attachment site on the hair follicle. There is no difference in the niche position between WT (grey) and C8cKO skin (red). **(E)** Probability plot of HFSC localization measured along hair follicles in WT (grey line) and C8cKO skin (red line). Pink shaded box marks the region where the probability is significantly different (p <0.05) between WT and C8cKO, n = 3 mice for each genotype (see supplementary table 3 for sample size of each set).”n.s.” indicates “not significant”.

**Figure 1: Figure Supplement 3.**
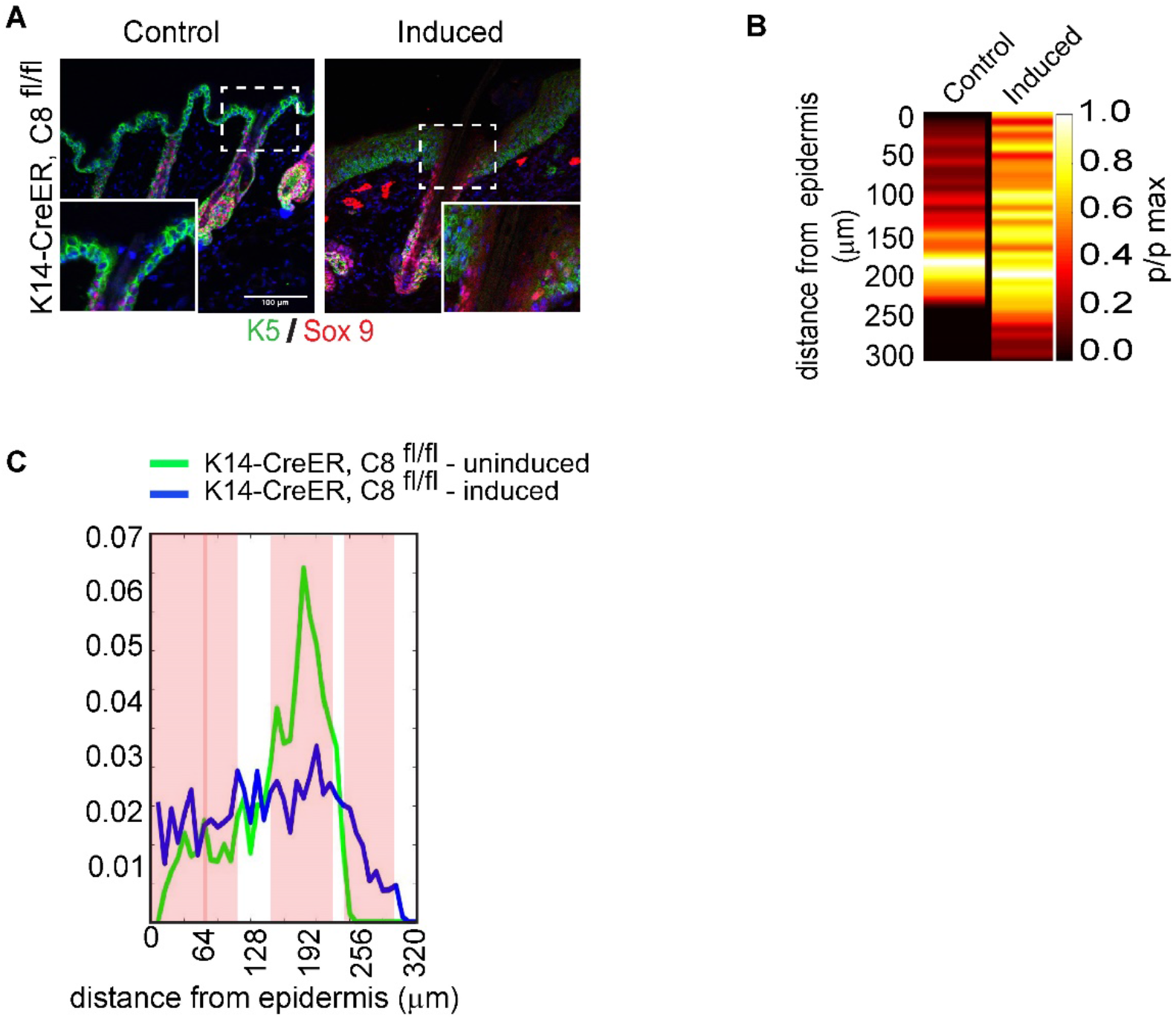
**(A)** Immunofluorescence images of HFSC migration to the epidermis is observed in tamoxifen inducible epidermal Caspase-8 knock-out adult mice compared to vehicle injected control. (**B**) Probability distribution of HFSCs along the hair follicle in tamoxifen inducible epidermal Caspase-8 knock-out adult mice compared to vehicle injected control, represented as a heat-map with distance from the epidermis represented on the y-axis. (**C**) Probability plot of HFSC localization measured along hair follicles in vehicle control injected (grey line) and tamoxifen injected inducible C8cKO skin (red line). Pink shaded box marks the region where the probability is significantly different (p <0.05), n = 3 mice for each conditioin (see supplementary table 4 for sample size of each set).

**Figure 1: Figure Supplement 4:**
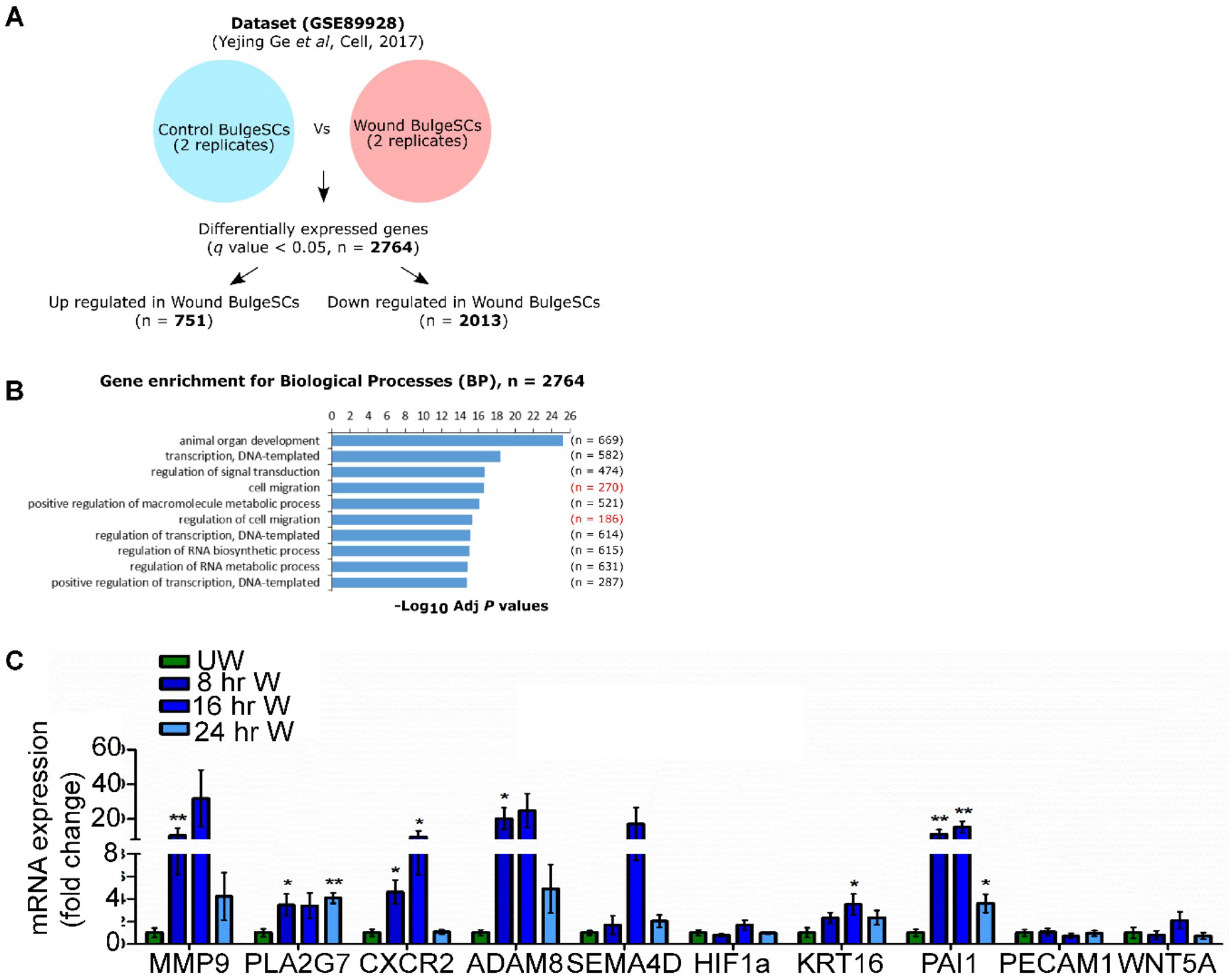
**(A)** Analysis of published RNA-seq data from homeostatic and wound-activated HFSCs collected at day 7 post wounding revealed a large number of differentially regulated genes. **(B**) Genes associated with cell migration form a major group of differentially regulated genes. Select genes from these categories were picked for further validation in our experiments. (**C**) Expression analysis of select migration associated genes at 8 (n=6), 16 (n=4) and 24 hours (n=4) of wound-healing in comparison to unwounded skin (green). Data is represented as the mean and S.E.M of the fold change relative to unwounded levels, “n” indicates the biological replicates and each biological replicate is a composite of three technical replicates. “*” indicates p<0.05, “**” indicates p<0.01.

**Figure 2: Supplementary Figure 1.**
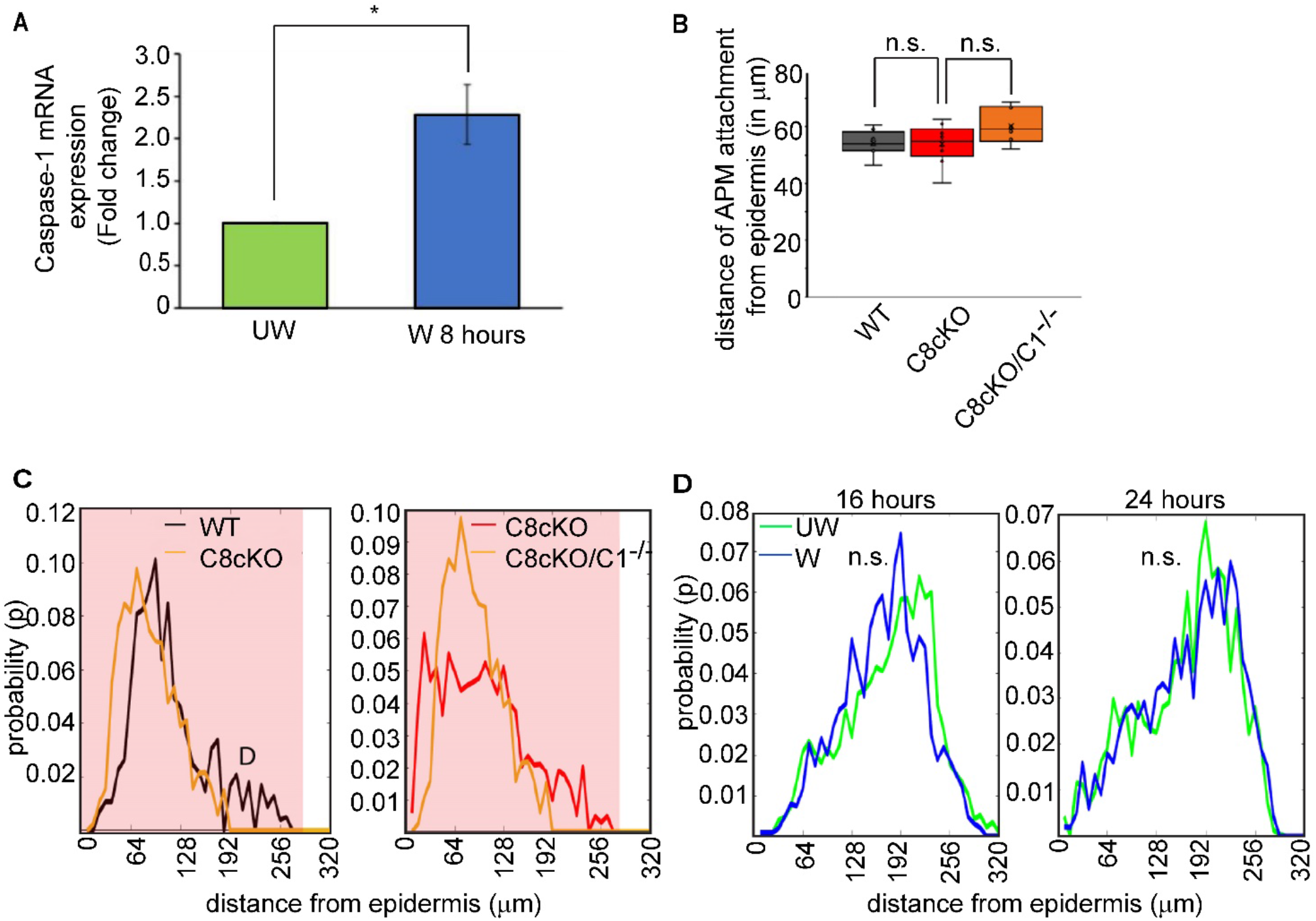
**(A)** Expression of Caspase-1 mRNA in unwounded (UW; green) and wounded (W) skin 8 hours post wounding (blue). Data represents mean and SEM from n= 4 mice. **(B)** In postnatal day 4 (p4) pups’ skin, the position of the HFSCs is determined by the distance of the APM attachment site on the hair follicle. There is no difference in the niche position between WT (grey), C8cKO (red), and C8cKO/C1^-/-^ (orange) skin. **(C)** Comparison of probability of HFSC localization measured along hair follicles in WT (black line) and C8cKO (orange line), and also between C8cKO (red line) and C8cKO/C1^-/-^ (orange line) skin. Pink shaded box marks the region where the probability is significantly different (p <0.05) between the represented pairs. The C8cKO/C1^-/-^ shows a rescue in phenotype towards WT distribution (see supplementary table 3 for sample size of each set). **(D)** Probability plot of HFSC localization in the C1^-/-^ skin measured along hair follicles in UW (green line) and W (blue line) skin 16 and 24 hours of wound-healing (see supplementary table 1 for sample size of each set). “n” indicates the biological replicate. “*” indicates p<0.05, “n.s.” indicates not significant. Note: Some of the WT and C8cKO mice chosen for analysis in this figure are same as those used for Figure 1 figure supplement 2F.

**Figure 2: Supplementary Figure 2.**
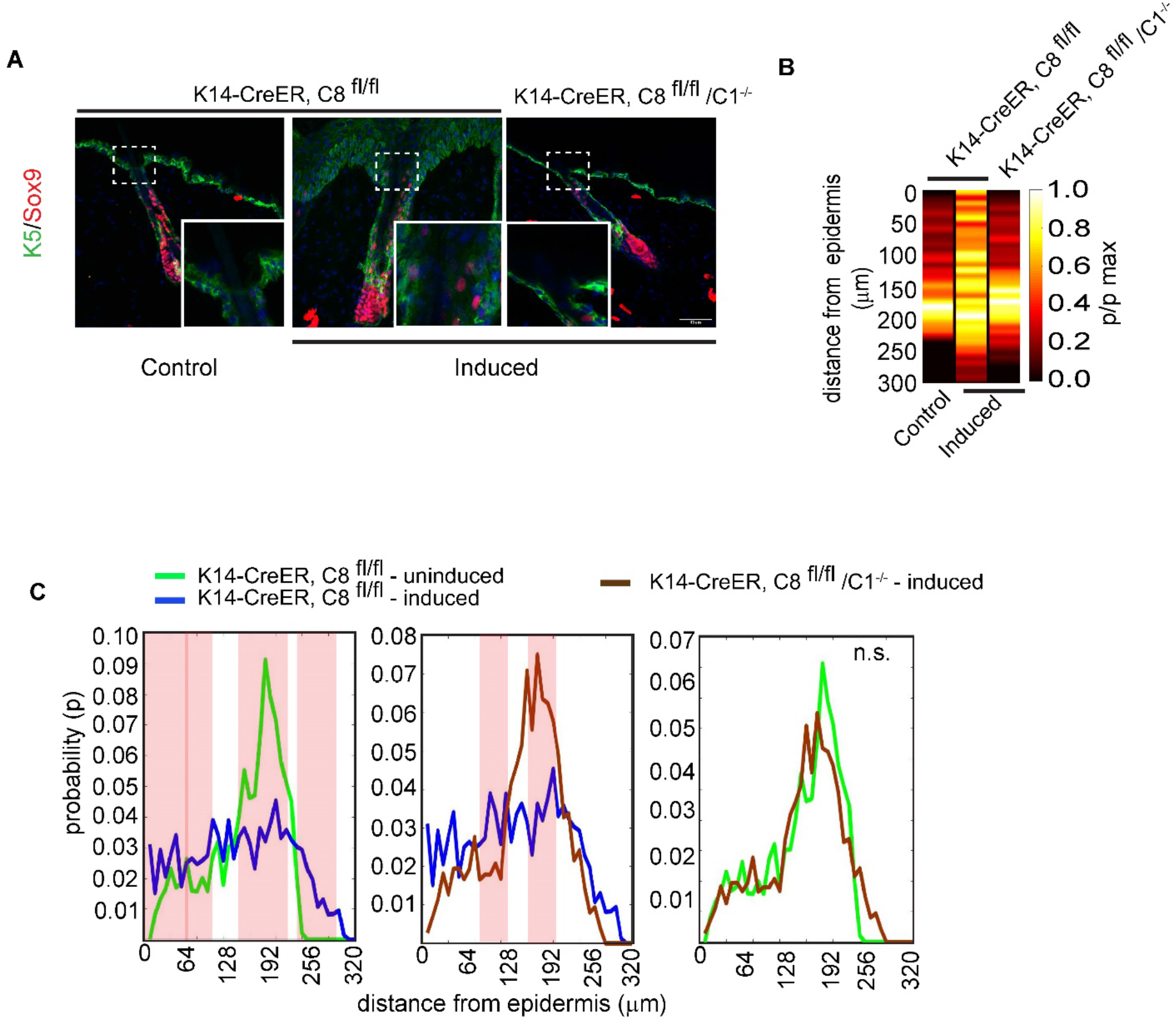
**(A)** Immunofluorescence images of localization of HFSCs (Sox9, red) in adult tamoxifen uninduced K14-CreER, C8^fl/fl^ mice, induced K14-CreER, C8 ^fl/fl^ mice and induced K14-CreER, C8 ^fl/fl^ mice in Caspase 1 null background (K14-CreER, C8 ^fl/fl^/C1^-/-^). Epidermis and hair follicles are marked by K5 (green). Insets show the infundibulum of representative hair follicles. (**B**) Probability distribution of HFSCs along hair follicles analyzed from adult uninduced K14-CreER, C8 ^fl/fl^ mice, induced K14-CreER, C8 ^fl/fl^ mice and induced K14-CreER, C8 ^fl/fl^ mice in caspase 1 null background. **(C)** Comparison of probability of HFSC localization measured along hair follicles in adult uninduced K14-CreER, C8 ^fl/fl^ mice, induced K14-CreER, C8 ^fl/fl^ mice and induced K14-CreER, C8 mice in caspase-1 null background. Pink shaded box marks the region where the probability is significantly different (p <0.05) between the represented pairs (see supplementary table 1 for sample size of each set). The induced K14-CreER, C8^fl/fl^ mice in caspase-1 null background shows a rescue in phenotype towards induced K14-CreER, C8 ^fl/fl^ mice with regards to the HFSC distribution “n” indicates the biological replicate. “*” indicates p<0.05, “n.s.” indicates “not significant”. Note: Induced and uninduced K14-CreER, C8^fl/fl^ mice chosen for analysis in this figure are same as those used for Figure 1 figure supplement 3.

**Figure 2: Supplementary figure 3.**
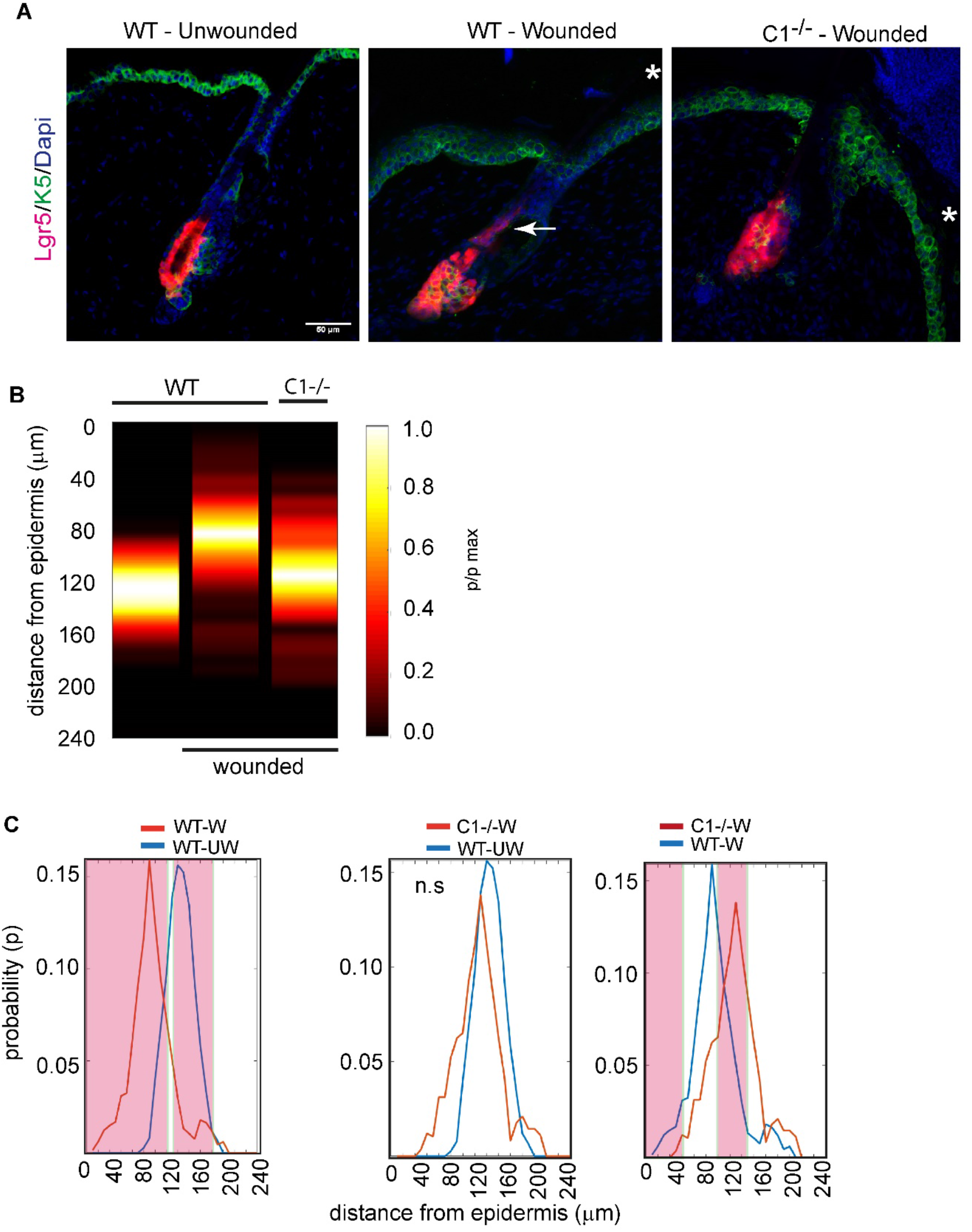
**(A)** Localization of HFSCs (Lgr5-tdT, red) in WT-Unwounded, WT-Wounded and Caspase-1 Null-Wounded skin. Epidermis and hair follicles are marked by K5 (green). The arrow points the migrated HFSC. “*” indicates the wound edge (**B**) Probability distribution of HFSCs along hair follicles analyzed from WT-Unwounded, WT-Wounded and Caspase-1 Null-Wounded skin, represented as a heat-map with distance from the epidermis represented on the y-axis. **(C)** Comparison probability of HFSC localization in wild type unwounded (WT-UW), wild type wounded (WT-W), and Caspase-1 null wounded (C1-/- W) skins. Pink shaded boxes marks the region where the probability is significantly different (p<0.05) between the presented pairs, at least n = 3 for each condition (see supplementary table 6 for sample size of each set).“n.s” indicates “not significant”.

**Figure 2: Supplementary Figure 4.**
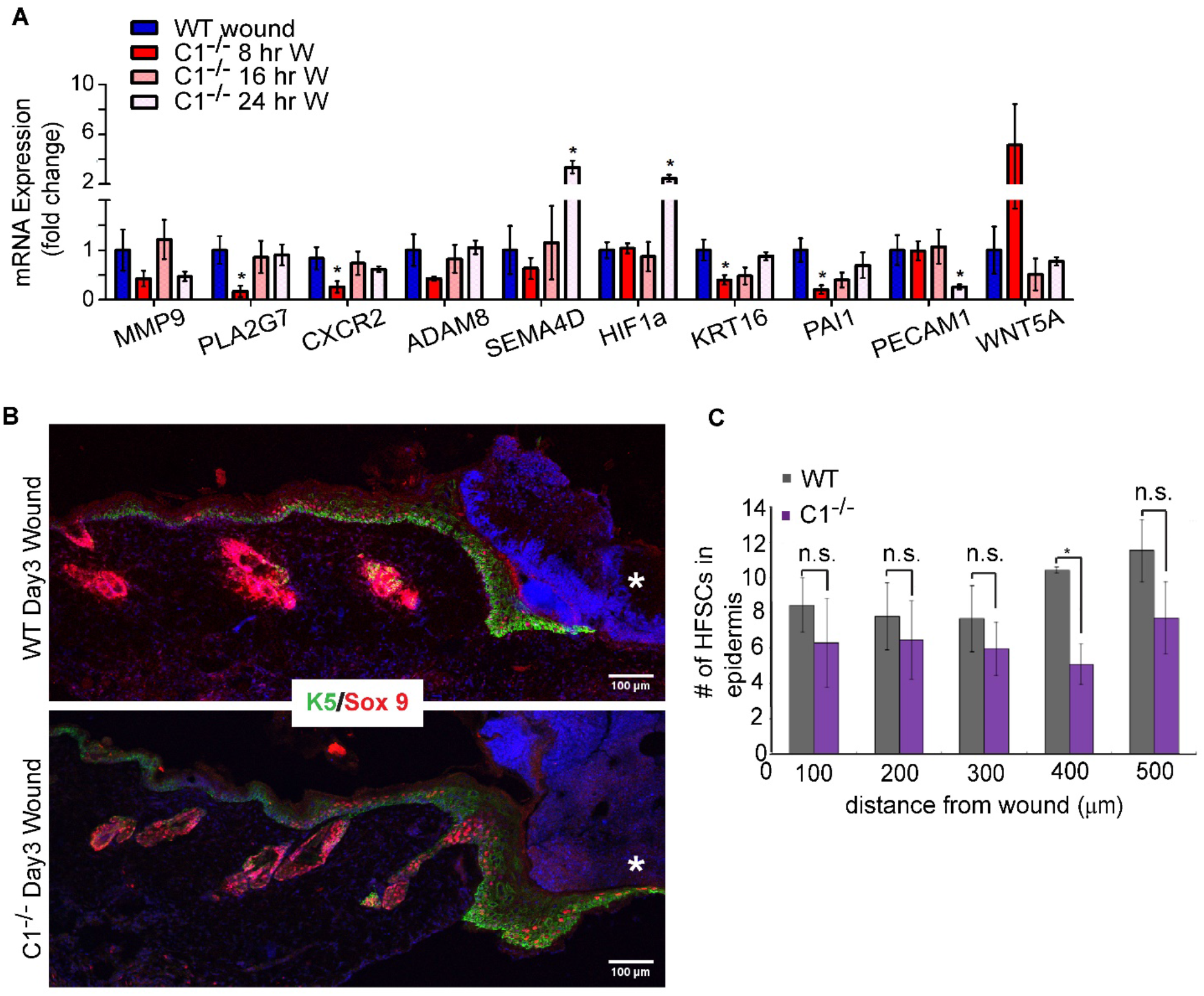
**(A)** Expression analysis of select migration associated genes in wounded skin of C1^-/-^ mice at B (n= 4) 16 (n=4) and 24 (n=3) hours (shades of pink) post wounding compared to wounded skin from wild-type mice normalized to 1(blue) Also, for each biological replicate, three technical replicates were performed. Data represented as mean and S.E.M. **(B)** Sox9 (red) and keratin 5 (K5, green) visualized in wound proximal skin 3 days post wounding in both WT and C1^-/-^ mice. “*” marks the wound edge. **(C)** Quantification of migrated HFSCs into the epithelial lip at 3 days post wounding reveals no difference between WT and C1^-/-^ wounds, n = 3 for each genotype. “n” indicates the biological replicates. “*” indicates p<0.05 “n.s.” indicates not significant.

**Figure 3: Supplementary Figure 1.**
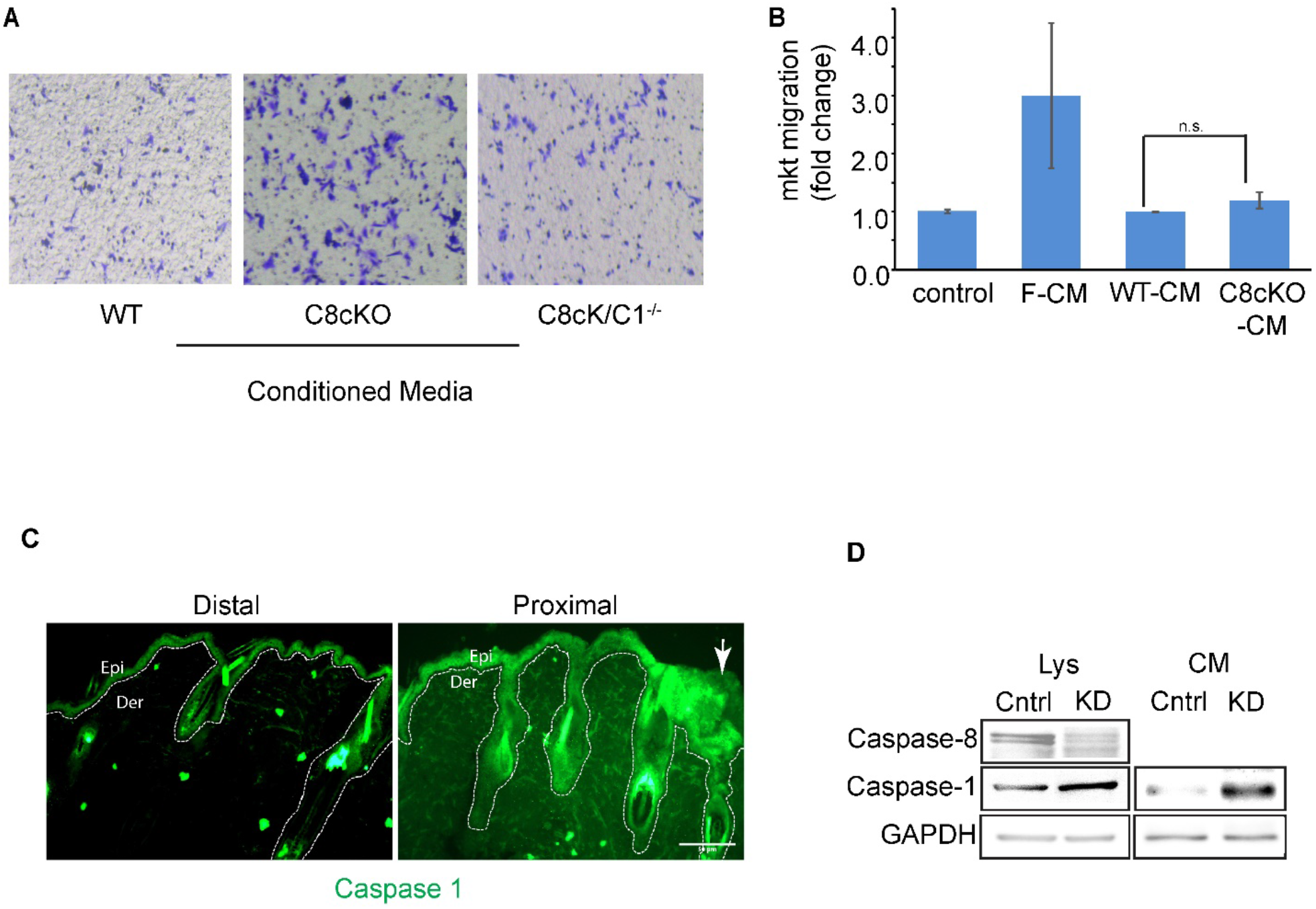
**(A)** HFSC migration in a Boyden chamber/transwell migration assay with conditioned media from WT epidermis (WT), C8cKO epidermis (C8cKO) and C8cKO-Caspase-1 null epidermis (C8cKO/C1^-/-^) in comparison to that from wild-type epidermis (WT, gray), Representive image of 4 biological replicates.(**B**) Migration of primary mouse epidermal keratinocytes measured by Boyden chamber assay in response to wild-type (WT-CM, gray) or caspase-8 conditional knockout epidermal conditioned media (C8cKO-CM, red). Feeder cell conditioned media (F-CM, green) is used as a positive control for inducing migration. Basal migration of cells with no chemoattractants is represented in yellow. Data shows mean and S.E.M. of of three biological replicates. **(C)** Wounding leads to secretion of Caspase-1 into the extracellular matrix. Compared to the wound distal area, a higher level of Caspase-1 can be observed proximal to the wound. Immunostaining against Caspase-1 is done on the unpermeabilized wounded mouse skin sections and representative image of three independent expreriments are presented. Brightness and contrast of the images have been adjusted equally. Distal (>3mm) and proximal (<1 mm) refer to the distance from the wound site. Arrow denotes the wound site and the dotted line marks the epidermis. Epi is epidermis and Der is dermis. **(D)** Detection of Caspase-1 in the lysate (Lys) and conditioned media (CM) from human epidermal keratinocytes (HaCat cells) transduced with scrambled shRNA, or shRNA against caspase-8 in knock-down cells (KD). Caspase-8 is probed in both cell lysates, which shows > 80% reduction in protein level, “n.s.” indicates “not significant”.

**Figure 4: Supplementary Figure 1.**
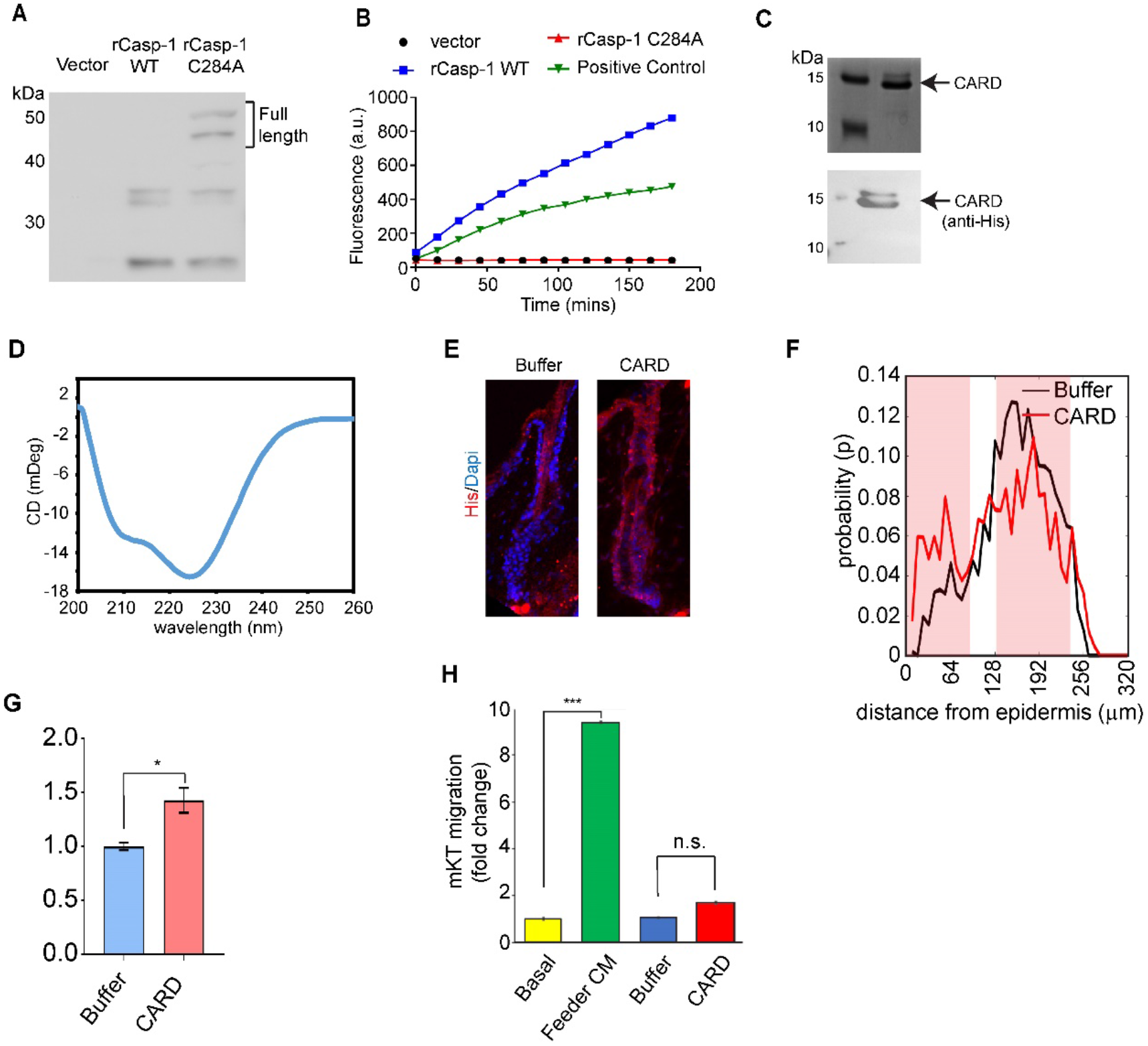
**(A)** Expression of 6x-His tagged wild-type (rCasp1-WT) and catalytically dead (rCasp1-C284A) recombinant murine caspase-1 proteins and vector expression proteins detected using an anti-Caspase-1 antibody. (**B**) Measurement of caspase-1 catalytic activity in a substrate cleavage assay of both the recombinant WT and C284A caspase-1 proteins in comparison to lysate obtained from HaCat cells (Positive control). Increase in fluorescence upon substrate (Ac-YVAD-AMC) cleavage is plotted on the y-axis. (**C**) Purified recombinant murine caspase-1 N-terminal domain CARD with C-terminal 10x-His tag is visualized by both silver stain (top panel) and by western blot using anti-His antibody (bottom panel). (**D**) CD spectra of recombinant Caspase-1 CARD. (E) Binding of CARD onto hair follicles visualized by anti-His antibody (red). (**F**) Probability plot of HFSC localization measured along hair follicles upon application of recombinant CARD and buffer on Caspase-1-/- skin. Pink shaded box marks the region where the probability is significantly different (p <0.05) between buffer and CARD, n = 3 mice for each condition (see supplementary table 5 for sample size of each set). **(G)** Analysis of caspase-1 null HFSC chemotaxis to recombinant CARD in comparison to buffer control from four biological replicates. Data shows mean and S.E.M. for both the experiments. (**H**) Transwell migration of primary mouse epidermal keratinocytes (mKT) in response to feeder conditioned media (green), buffer (blue) and CARD (red). Basal level of migration is shown in yellow. Data is composite of three biological replicates.“**8” indicates p<0.001, “n.s.” indicates not significant.

**Figure 4: Supplementary figure 2.**
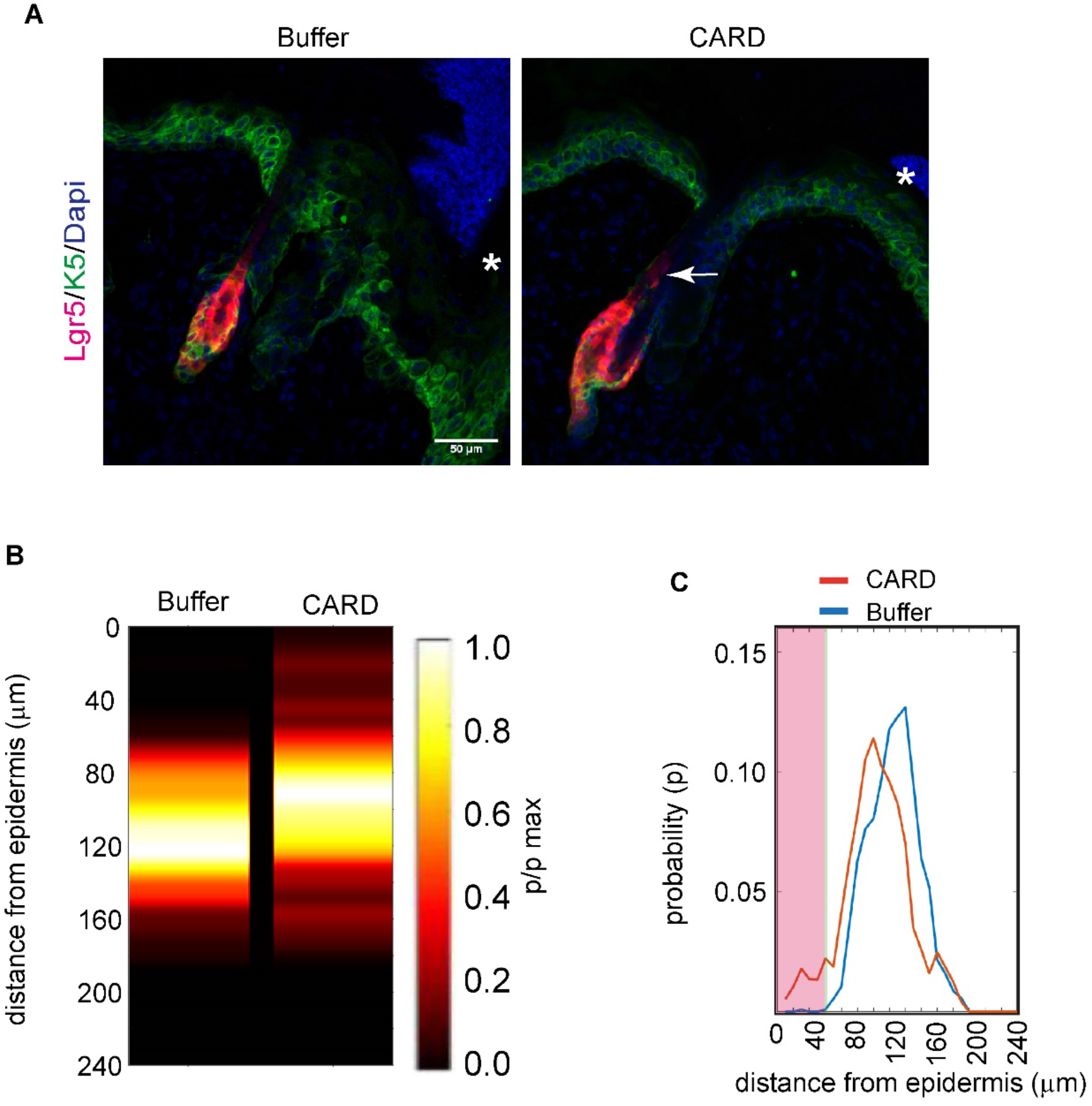
**(A)** Representative images showing the migration of HFSCs towards the epidermis upon application of recombinant CARD or buffer on the wounded caspase-1 Null skin. The arrow points the migrated Lgr5 makrked HFSC. “*” marks the wound edge. (**B**) Probability distribution of HFSCs in wound-proximal hair follicle from buffer of CARD treated skins. **(C)** Comparision probability of HFSC localisation in wild type unwounded (WT-UW), wild type wounded (WT-W) and caspase-1 null wounded (C1-/- W) skins. Pink shaded boxes marks the region where the probability is significantly different (p<0.05) between the presented pair, n = 3 for each condition (see supplementary table 6 for sample size of each set).

**Supplementary table 1:** Animal and hair follicle numbers used in statistical analysis of HFSC distribution in adult WT and Caspase-1 null mice unwounded (UW) and wounded (W) skin. (For Figure 1 A and B, Figure 2 C and D)

| Condition | WT |  | Caspase-1 null |  |
| --- | --- | --- | --- | --- |
|  | Animal number (n) | Number of hair follicles from each animal | Animal number (n) | Number of hair follicles from each animal |
| 8 hours UW | 1 | 1 |  |  |
|  | 2 | 2 |  |  |
|  | 3 | 5 |  |  |
|  | 4 | 3 |  |  |
|  | 5 | 7 |  |  |
| 8 hours W | 1 | 1 |  |  |
|  | 2 | 3 |  |  |
|  | 3 | 3 |  |  |
|  | 4 | 7 |  |  |
| 16 hours UW | 1 | 8 | 1 | 10 |
|  | 2 | 9 | 2 | 3 |
|  | 3 | 5 | 3 | 3 |
|  | 4 | 7 | 4 | 3 |
|  | 5 | 5 | 5 | 5 |
|  |  |  | 6 | 3 |
| 16 hours W | 1 | 6 | 1 | 2 |
|  | 2 | 17 | 2 | 2 |
|  | 3 | 7 | 3 | 4 |
|  |  |  | 4 | 5 |
| 24 hours UW | 1 | 4 | 1 | 4 |
|  | 2 | 6 | 2 | 9 |
|  | 3 | 4 | 3 | 4 |
| 24 hours W | 1 | 5 | 1 | 2 |
|  | 2 | 6 | 2 | 7 |
|  | 3 | 3 | 3 | 6 |

**Supplementary table 2:** Animal and hair follicle numbers used in statistical analysis of Lgr5 tdT HFSC distribution in lineage tracing experiments. (For Figure 2: Figure Supplement 3)

|  | WT |  | Caspase-1 null |  |
| --- | --- | --- | --- | --- |
| Condition | Animal number (n) | Number of hair follicles from each animal | Animal number (n) | Number of hair follicles from each animal |
| Unwounded | 1 | 3 |  |  |
|  | 2 | 3 |  |  |
|  | 3 | 3 |  |  |
| Wounded | 1 | 2 | 1 | 3 |
|  | 2 | 3 | 2 | 3 |
|  | 3 | 3 | 3 | 4 |
|  | 4 | 4 | 4 | 1 |

**Supplementary table 3:** Animal and hair follicle numbers used in statistical analysis of HFSC distribution in p4 mice. (For Figure 1 C and D, Figure 2 A and B)

| Genotype | Animal number (n) | Number of hair follicles from each animal |
| --- | --- | --- |
| WT | 1 | 5 |
|  | 2 | 6 |
|  | 3 | 6 |
| C8cKO | 1 | 5 |
|  | 2 | 6 |
|  | 3 | 6 |
| C8cKO/C1<sup>-/-</sup> | 1 | 4 |
|  | 2 | 7 |
|  | 3 | 7 |

**Supplementary table 4:** Animal and hair follicle numbers used in statistical analysis of HFSC distribution in adult inducible C8cKO mice. (For Figure 1: Figure Supplement 3 and Figure 2: Supplementary Figure 2)

| Genotype | Animal number (n) | Nmber of hair follicles from each animal |
| --- | --- | --- |
| K14-CreER, C8<sup>fl/fl</sup> uninduced | 1 | 3 |
|  | 2 | 3 |
|  | 3 | 3 |
| K14-CreER, C8<sup>fl/fl</sup> induced | 1 | 4 |
|  | 2 | 3 |
|  | 3 | 3 |
| K14-CreER, C8<sup>fl/fl</sup> / C1<sup>-/-</sup> induced | 1 | 3 |
|  | 2 | 3 |
|  | 3 | 3 |

**Supplementary table 5:** Animal and hair follicle numbers used in statistical analysis of HFSC distribution in wounded Caspase-1 mice upon CARD application. (For Figure 4 E and F)

|  | Caspase-1 null |  |
| --- | --- | --- |
| Condition | Animal number (n) | Number of hair follicles from each animal |
| Wounded+Buffer | 1 | 2 |
|  | 2 | 4 |
|  | 3 | 1 |
|  | 4 | 2 |
| Wounded+CARD | 1 | 3 |
|  | 2 | 3 |
|  | 3 | 2 |
|  | 4 | 2 |

**Supplementary table 6:** Animal and hair follicle numbers used in statistical analysis of Lgr5 tdT HFSC distribution in lineage tracing experiments upon CARD application. (For Figure 4: Supplementary figure 2)

|  | Caspase-1 null |  |
| --- | --- | --- |
| Condition | Animal number (n) | Number of hair follicles from each animal |
| Wounded+Buffer | 1 | 8 |
|  | 2 | 5 |
|  | 3 | 3 |
| Wounded+CARD | 1 | 7 |
|  | 2 | 4 |
|  | 3 | 3 |

**Supplementary Table 7:**
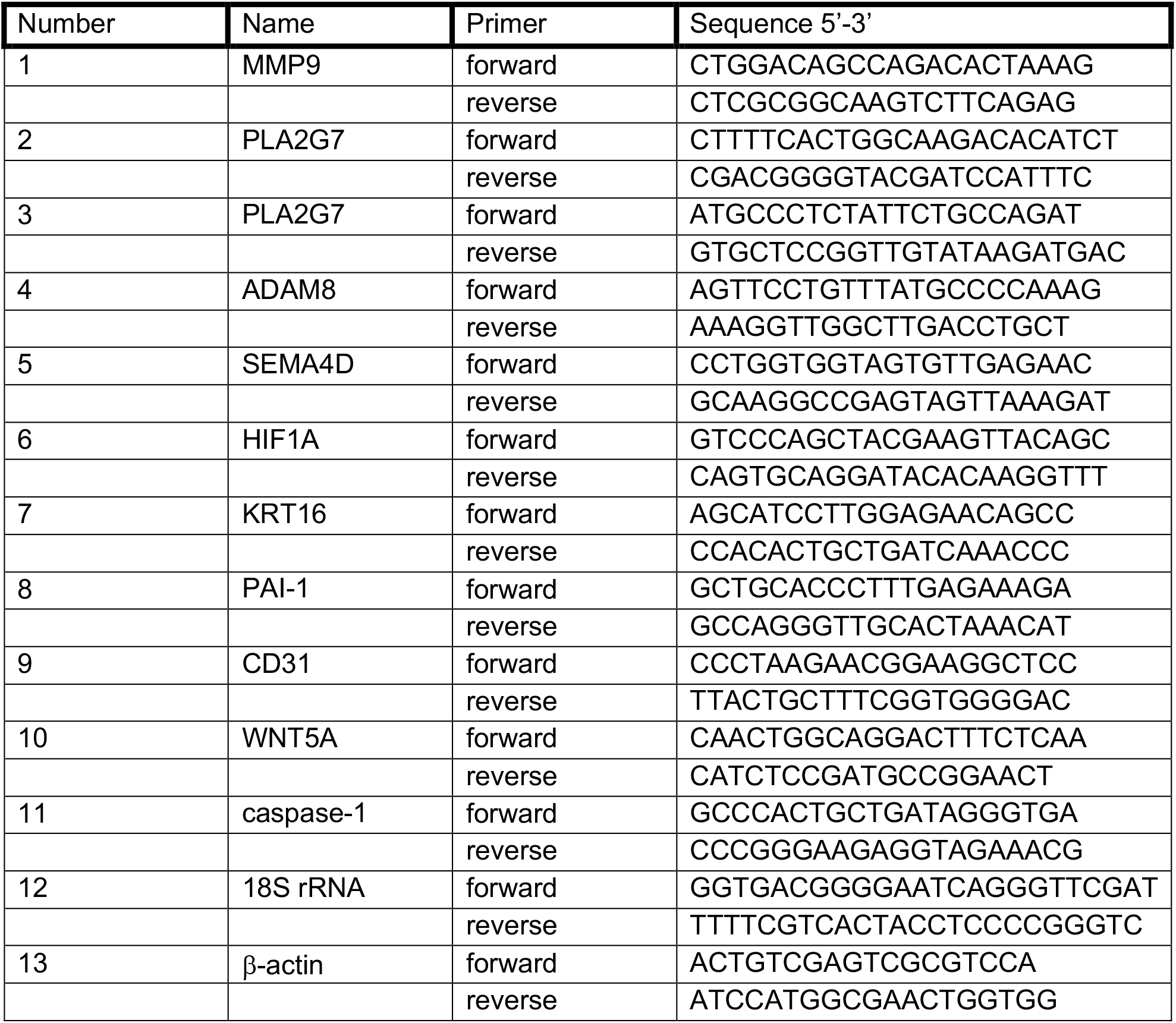
List of primers used for qPCR.

